# An Integrated Photoreceptor-to-RGC Stimulation Circuit for Intraocular Visual Prostheses

**DOI:** 10.64898/2026.05.15.725457

**Authors:** Vedika Bedi, Mujeeb U. Chaudhry

## Abstract

Visual prostheses face a critical miniaturisation challenge: converting photoreceptor signals to biologically appropriate retinal ganglion cell (RGC) stimulation patterns within the spatial constraints of intraocular implants. Existing systems rely on external microcontrollers for signal processing, limiting scalability for high-density pixel arrays. This paper presents an integrated per-pixel circuit architecture that directly converts photocurrent into frequency-modulated current pulses that match RGC activation thresholds.

The design targets are established through NEURON computational modelling of red-green colour-opponent midget RGCs, identifying stimulation thresholds of +0.1nA to +3.5nA for depolarisation and -0.1nA for repolarisation. The proposed circuit combines a transimpedance amplifier, a voltage-controlled oscillator with a Schmitt trigger, and a current-controlled output stage to generate biphasic pulses within these thresholds. A complementary output provides lateral inhibition, reducing crosstalk between adjacent RGC stimulation sites.

Photoreceptor integration is achieved using P3HT:PCBM organic photodiodes for cone-associated RGCs and phototransistors for rod-associated RGCs, validated through OghmaNano finite element simulations. The photodiode circuit produces output frequencies of 2.5Hz (dark) to 600Hz (100 *W/m*^2^), matching reported RGC response ranges. This architecture eliminates external processing requirements, enabling scalable high-density retinal prostheses design.

## I. Introduction

**T**HIS paper uses the architecture of the visual system to replicate signals and construct an artificial vision system for individuals with visual impairment. The three primary causes for blindness are; retinal damage, glaucoma (Retinal Ganglion Cell/Optic nerve damage), and neurological issues affecting the brain’s visual processing [1]. The most prevalent issues, retinal damage and glaucoma, can be both artificially stimulated and replicated.

Several types of visual prostheses have been proposed and classified by their location along the visual pathway, including subretinal, epiretinal and other cortical prostheses. Alpha IMS (Retina Implant AG; Reutlingen, Germany) is a subretinal implant with a 1500 photodiode light sensitive array [2] positioned in the layer of degenerated photoreceptors, stimulating the bipolar cells - the link between Retinal Ganglion Cells (RGCs) and the photoreceptors.

The biggest challenge with retinal implants is the miniaturisation of the electronic circuitry. Photodiode pixels have been successfully manufactured at the micrometre scale and produce an output photocurrent. However, packaging the remaining signal-conditioning electronics within the tight confines of the eye and retina is a challenge. Alpha IMS tackles this problem by positioning the flat “vision chip” with 1500 photodiodes, coupled via 1500 amplifiers directly to the stimulation electrodes.[2].

Although CurvIS (a hemispherically curved image sensor array) developers [3], have successfully developed an ultra-thin and soft optoelectronic pixel array matching the mechanical properties of the retina, they have not achieved electronic miniaturisation. The system involves vertically stacked electrodes and photo-transistors on the curved pixel array, with a pulse stimulating the retina. However, the photocurrent must travel to an external soft Flexible PCB (FPCB) for processing before returning to stimulate the electrodes. The photocurrent response is amplified by a transimpedance amplifier and an inverter, becoming the input to a micro-controller which measures the amplified signal, processes it and produces programmed electrical pulses [3] on an external soft FPCB. Although the use of microcontrollers simplifies the programming and delivery of pulse trains, scalability becomes an issue, as each microcontroller with a single CPU can only deliver a single pulse train for a single pixel at a time. With an array of over 1,000 pixels, this becomes an issue.

The optic nerve receives signals from approximately 125 million photo-receptor cells. It connects to RGCs, which bundle up with more than a million fibres [4], forming the base of the optic nerve and sending signals to the brain. RGCs, characterised by distinct morphologies, serve as the foundation for visual processing, with each group fulfilling specialised roles. Although mid-retinal implants such as Alpha IMS may facilitate natural information processing along the visual pathway, retinal damage often affects multiple cell layers simultaneously, limiting the effectiveness of approaches that rely on intact bipolar cell function. While some studies have investigated direct optic nerve stimulation [5], stimulating the nerve bundle directly does not replicate natural vision signalling as effectively as targeting individual RGCs [6]. Information transfer between neurons occurs at synapses, with Glutamate serving as the excitatory neurotransmitter and GABA as the inhibitory neurotransmitter [7].

This paper proposes an integrated per-pixel circuit architecture that emulates RGC nerve fibre responses at biologically appropriate potentials, aligning electrochemical signals with visual inputs to achieve selective activation. The design incorporates lateral inhibition between opponent RGC cell pairs to prevent crosstalk between neighbouring axons.

## II. RGC Stimulation Requirements

Investigations into colour-opponent cells highlight the pivotal role of midget ganglion cells at the foveal centre in conveying red-green colour signals to the brain [9]. This study focuses on advancing computational models using the NEURON Simulation Environment, leveraging extensive research [10] [8] [9] into the morphology and biophysical attributes of these cells. Understanding how RGCs respond to electrical stimulation and selectively targeting them based on these traits could significantly enhance the performance of current retinal prosthetic systems [11].

### A. Red-Green Colour Opponent Midget RGC Morphology

Neuronal cells can be categorised as ON cells (responsive to light increments), OFF cells (responsive to light decrements), and ON-OFF cells (responsive to both light increases and decreases) [12]. The fovea, crucial for high acuity vision, contains midget RGCs known for their small soma size [13]. They exhibit receptive fields with a centre-surround organisation, facilitating the encoding of red-green opponent signals [13]. Specifically, Red ON/Green OFF Cells respond to glutamate under red light and are inhibited by green light, causing increased firing rates in RGCs with cone cells under excitation of long wavelengths. Conversely, Green ON/Red OFF cells are excited by green light and inhibited by red, causing increased firing rates with cone cells under excitation of short wavelengths [13].

In Figure 2, note how the magnitude of the RGC response doesn’t vary; only the frequency of firing rates changes. This is how the neural system responds to an increase in stimuli.

### B. RGC Biophysics and Ion Kinetics

NEURON is a computational tool used in neuroscience to model the behaviour of individual neural cell types, neurons, and neural networks [15]. Mathematical models that mimic the electrical activity of neurons can be constructed by considering both the chemical mass and the electrical charge conservation laws. Action potentials result from the sum of currents entering the cell from neighbouring regions and externally injected currents such as synaptic inputs [15]. The short electrical pulses are fired across synapses when synaptic inputs exceed the cell’s built-in threshold value, a process mediated by the excitatory neurotransmitter Glutamate across RGC synapses [16].

The Hodgkin-Huxley (HH) Model provides a conductance-based representation of action potential generation, characterising three channels by resistance or conductance. The leakage channel maintains the resting membrane potential, while the sodium and potassium channels exhibit voltage-dependent conductance responsible for action potential generation. Sodium influx through activation gates leads to depolarisation, while slower-responding potassium gates contribute to repolarisation by outward flow of potassium ions [17]. GABA is the inhibitory neurotransmitter that activates these potassium channels, while Glutamate is the excitatory neurotransmitter.

This natural inhibitory mechanism inspires the lateral inhibition strategy employed in the proposed circuit to reduce cross-communication between closely packaged stimulation sites.

Action potentials, once initiated, maintain a constant amplitude and duration, regardless of stimulus strength. However, variations in stimulus intensity are encoded through changes in firing frequency (Fig. 2), with stronger stimuli producing higher pulse frequencies. Research shows this frequency range to vary from 20*Hz* to 700*Hz* depending on RGC cell types and stimulus intensity [11], establishing the target frequency range for the circuit design.

### C. Computational Modelling

The structural morphology of a red-green colour-opponent midget RGC cell was configured in NEURON 8.2, as shown in Figure 1, with biophysical properties detailed in the supplementary Table S1. The HH model was implemented at the soma, while a passive model represented leak conductances across dendritic extensions. To replicate neurotransmission from a bipolar cell to an RGC cell, an extracellular mechanism known as a current clamp (IClamp), was applied via the Alpha Synapse Tool with configurable delay, duration, and amplitude of current injection.

**Figure 1:**
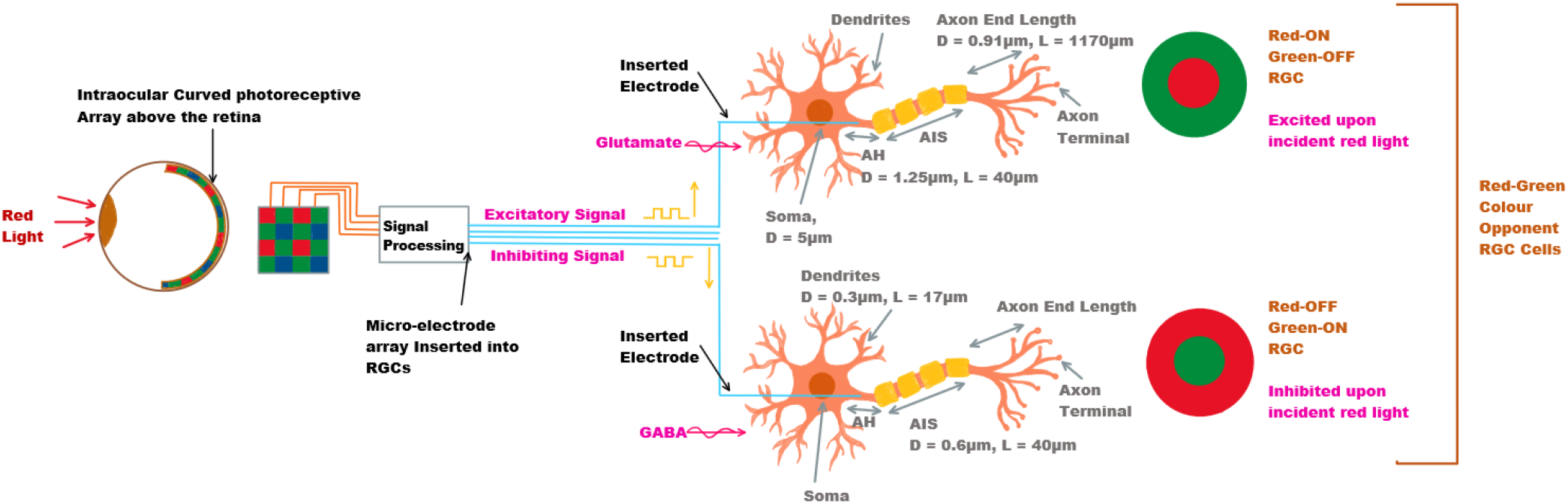
An illustration of the intraocular photoreceptive array inducing both excitatory and inhibitory signals in red-green colour opponent cells upon incident red light. For both, electrodes are shown to be inserted into the RGC cell axon and stimulated using a pulse via a function generator. Glutamate and GABA are illustrated to show synaptic neurotransmission under normal retinal operation. The neuron model shows the morphology of the RGC; where the soma is the cell body containing the nucleus, the dendrites receive inputs from other cells, and the axon carries the electrical impulse to be received by other neurons. [8].

**Figure 2:**
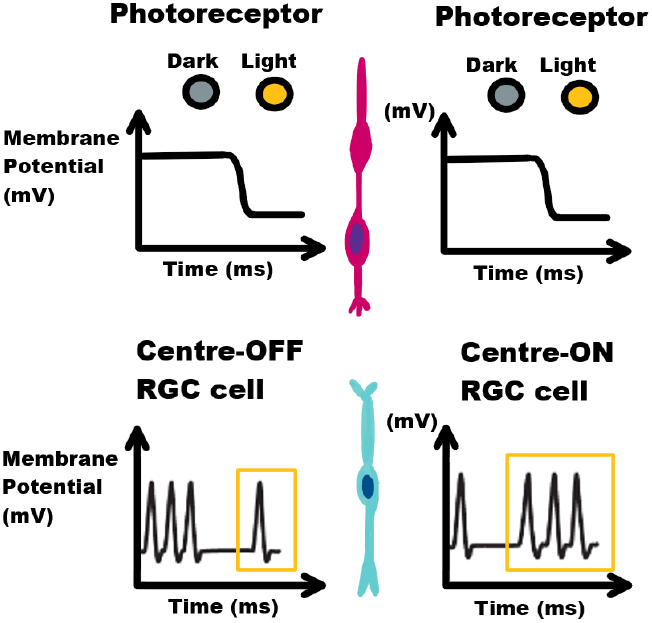
All photoreceptors respond with hyperpolarisation when transitioning from dark to illuminated conditions. Centre-OFF RGCs elevate their firing rates in darkness and diminish them in response to light stimuli. Conversely, Centre-ON RGCs increase their firing rates upon exposure to light and decrease them in darkness. [14]

Figure 3 shows the activated potential across the RGC section (soma, axon hillock, AIS, axon terminal) with varying current injection at the synaptic dendrite. Upon current injection at the dendrite, it induces a localised change in the membrane potential. The gradual drop in membrane potential from soma to axon terminal is attributed to passive propagation along the neuron’s cable properties. The simulation does not fully account for axon myelination, which would reduce these passive leakages in biological cells.

**Figure 3:**
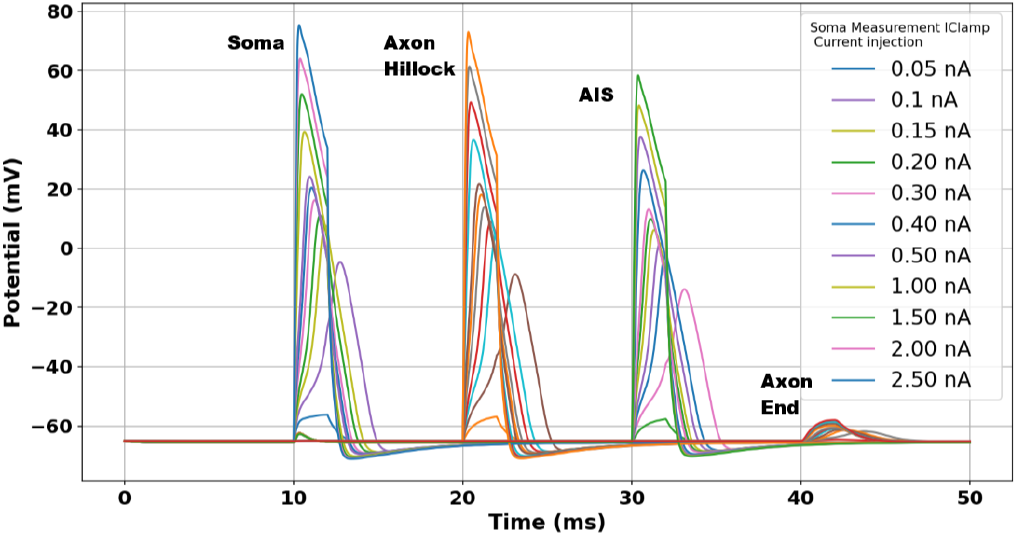
This graph from NEURON simulations shows the potential at the Soma, Axon Hillock, AIS, and Axon End, each with a varying current injection from 0.05 nA to 2.50 nA

Figure 4 reveals that increasing pulse duration produces an apparent “saturation” effect, attributable to excessive depolarisation. A 1-2ms current pulse shows a characteristic spike followed by rapid decline, indicating the HH-modelled potassium ion release for repolarisation. However, this natural repolarisation mechanism proves insufficient for longer current pulses, establishing the need for an active inhibiting signal in the artificial stimulation circuit.

**Figure 4:**
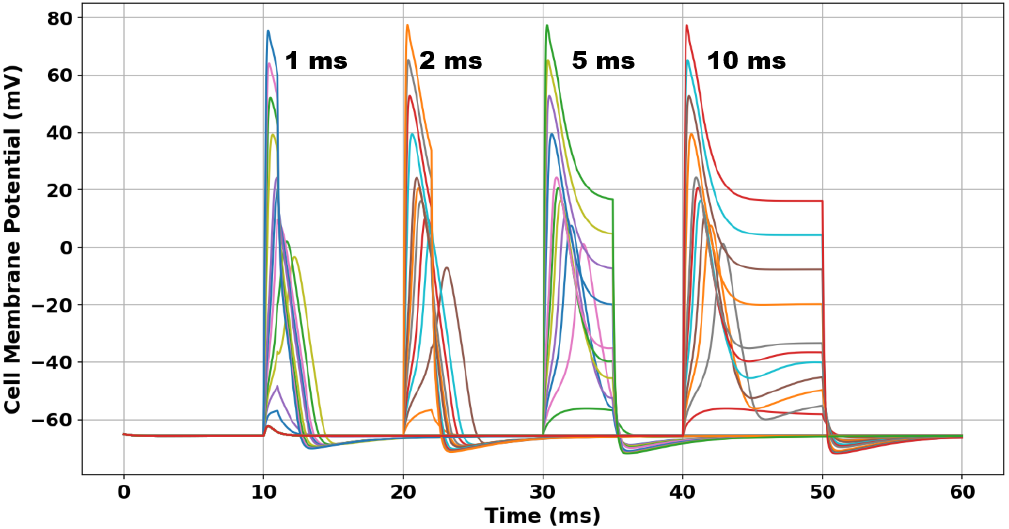
Graph from NEURON simulations shows the soma membrane potential and current injection but with varying levels of pulse duration (1ms, 2ms, 5ms, 10ms).

Biphasic stimulation pulses are therefore essential (Fig. 5). The cathodic phase depolarises the membrane and elicits a neural response, while a subsequent anodic phase discharges the accumulated charge around the electrode. Although some studies employ an equal anodic and cathodic current injection, NEURON results and proposed neuron circuit models such as the Izhikevich model [18], suggest a slow repolarisation aiming to prevent intense hyperpolarisation.

**Figure 5:**
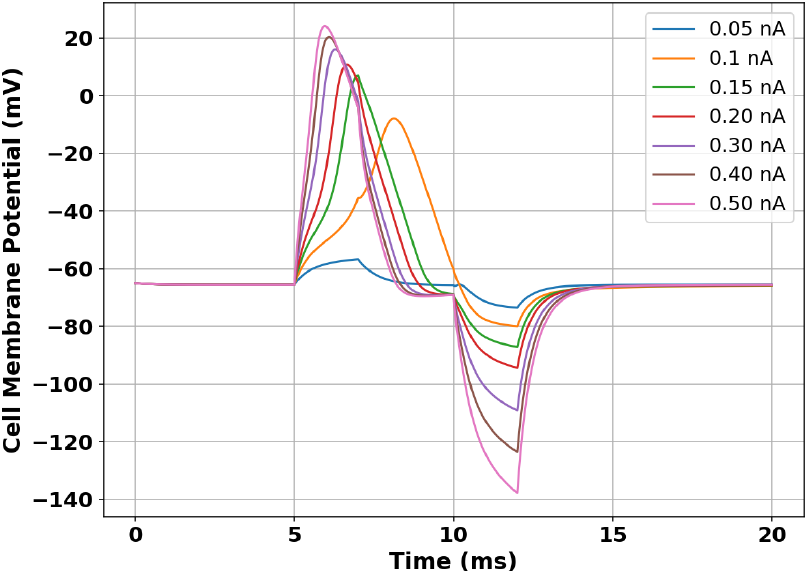
NEURON results from a biphasic pulse current injection. The first 2ms phase of the pulse (cathodic current) depolarises the cell membrane and elicits a natural response. The second 2ms phase (anodic current) balances the first phase to prevent net charge accumulating on the electrode. An intraphase delay of 2ms separates the pulses so that no reversal of the physiological effects occur [19].

Table I summarises the membrane potential thresholds and the equivalent current pulses, establishing the design parameters for the proposed circuit. Depolarisation begins to occur at a threshold of approximately -55mV (Fig. 3) [8]. The neuron completely depolarises at a membrane potential of +80mV once the sodium channels are fully open, requiring a cathodic pulse of 2.5nA when artificially stimulated. For repolarisation, however, an anodic pulse of -0.1nA is sufficient (Fig. 5).

**Table I.**
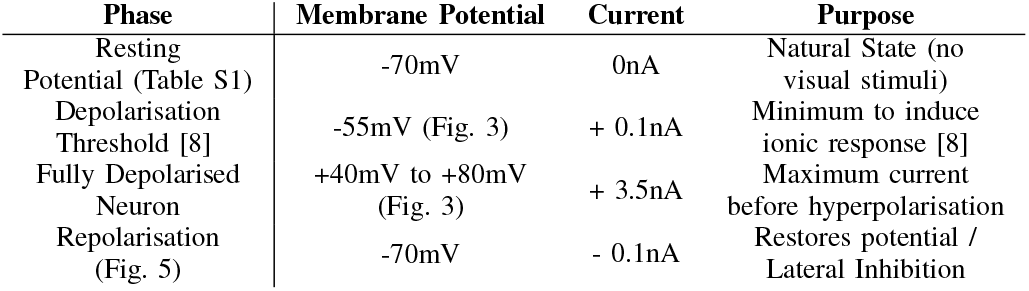
Current Pulse Requirements.

These four parameters — minimum depolarisation current (+0.1nA), maximum depolarisation current (+2.5nA), repolarisation current (−0.1nA), and frequency range (20-700Hz) — form the specifications for the stimulation circuit presented in Section III.

In conventional camera systems, digital processing algorithms handle demosaicing, colour extraction, and noise reduction. In the proposed artificial retina, these functions are simplified by directly stimulating RGC cells at frequencies corresponding to different light wavelengths — the brain’s visual cortex naturally handles colour perception and adaptation to lighting conditions. However, lateral inhibition remains necessary within the photoreceptive array to elevate signal-to-noise ratio and improve edge contrast, motivating the complementary output stage in the proposed circuit (Section III).

## III. Signal Processing and Neuronic Circuitry Development

### A. Proposed Circuit

The circuit presented in this paper attempts to provide a biologically realistic pulse train based on the photo-response of RGC cells. The circuit design (Fig. 6) aligns with the threshold parameters necessary for RGC activation, based on the NEURON computational modelling results. See the supplementary information (Table S2 and S3) for a breakdown of circuit parameters and FET aspect ratios.

**Figure 6:**
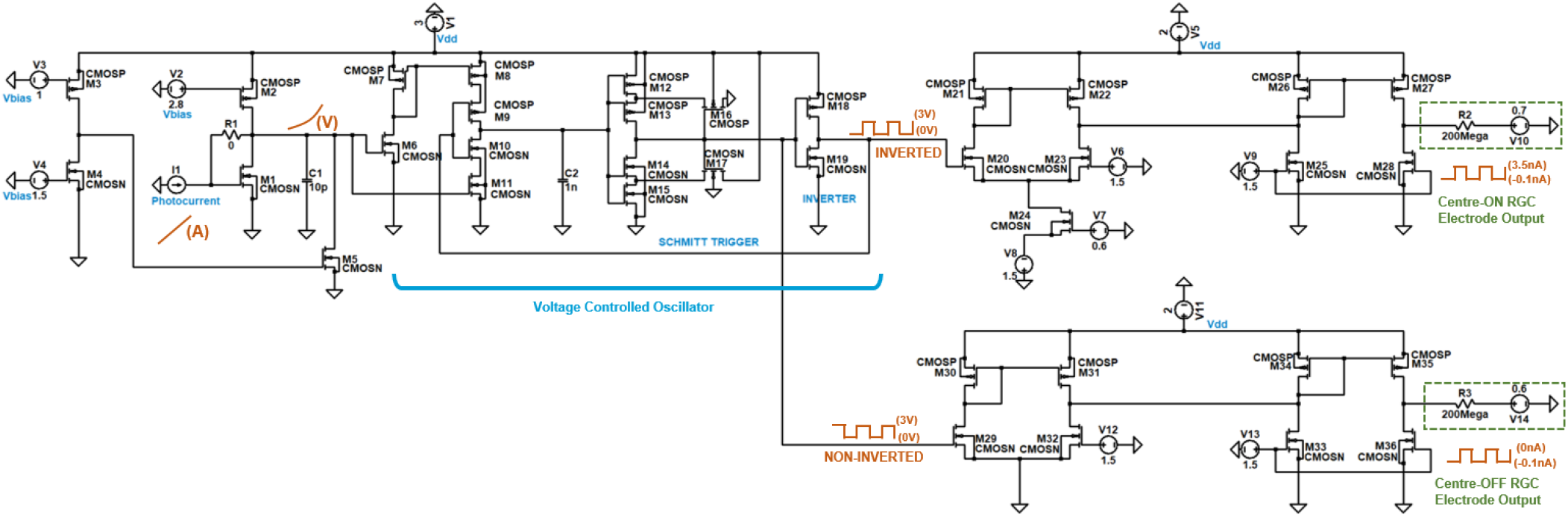
Proposed neuron spiking circuit (enlarged in Fig. S1) that converts a photocurrent to a suitable output current meeting the threshold requirements for RGCs. The electrodes (R2 and R3) induce an electrochemical response in the RGCs upon ‘firing’.

#### 1) Transimpedance Amplifier

Transistors M1 to M5 are the initial part of the circuit where the photoreceptor generates a current proportional to the light intensity, which is then converted to a voltage (M1 and M2). A current sink is created by M3, M4 and M5, ensuring that in the absence of photocurrent, the resulting voltage output is minimal and insufficient to influence the subsequent stages of the circuit. C1 smooths the photo-voltage output.

#### 2) Schmitt Trigger - Pulse Generation

The photo-voltage that is now proportional to light intensity, is input to a current mirror and CMOS circuit (M6 to M11) that provides biasing and stability to the Schmitt trigger and the output CMOS inverter (M18 and M19). The Schmitt Trigger is realised through a combination of P-channel and N-channel MOSFETs (M12 to M17) which provide the necessary hysteresis through positive feedback. The capacitor C2 is connected to the input of the Schmitt Trigger. It charges and discharges between the thresholds set by the Schmitt Trigger, thus creating an oscillation. The rate of charging and discharging, and therefore the frequency of the oscillation, is determined by the size of C2. Meeting neuron firing rate requirements and optimising the circuit meant that C2 was chosen to be between 0.1nF - 1nF depending on cell type (Fig. 13). Essentially, this circuit is a voltage controlled oscillator (VCO) and the magnitude of the output following the inverter, is a pulse train from 0V to 3V.

#### 3) Current Controlled Amplifier - Primary Current Output

A differential amplifier (M20 to M24) shifts the 0V to 3V Schmitt Trigger pulse train into a -170mV to +100mV pulse train. A current mirror (M25 to M28) takes the reference current created by the differential amplifier and reduces it down to a more appropriate level using different MOSFET aspect ratios (W/L). Lastly, R2 represents the electrode that induces the pulse in the Centre-ON RGC cell. The high resistance electrode is necessary for maintaining the current within the nanoampere (nA) range, aligning with the current levels observed in NEURON simulations.

#### 4) Lateral Inhibition - Complementary current Output

The electronics within this segment of the circuit are designed to mirror the primary current output, albeit with different MOSFET aspect ratios. This adjustment is purposefully executed to generate pulses ranging from 0A to -0.1nA to achieve the desired current modulation.

In this instance, the input to this circuit (M29 to M36) is taken from the non-inverting output of the Schmitt trigger, generating a complementary output. This mechanism facilitates lateral inhibition among the adjacent RGCs, effectively mitigating signal interference, or “cross-talk.” Consequently, when the primary current output registers a high level (+3nA), the corresponding complementary current is produced at a low level (−0.1nA). Conversely, when the primary current is at a low level (−0.1nA), indicative of the depolarisation of an RGC, the complementary current is then output at a high level (0A). This lateral inhibition system operates as seen in Figure 7 and utilises pixel pairs based on RGC opponent cell pairs.

**Figure 7:**
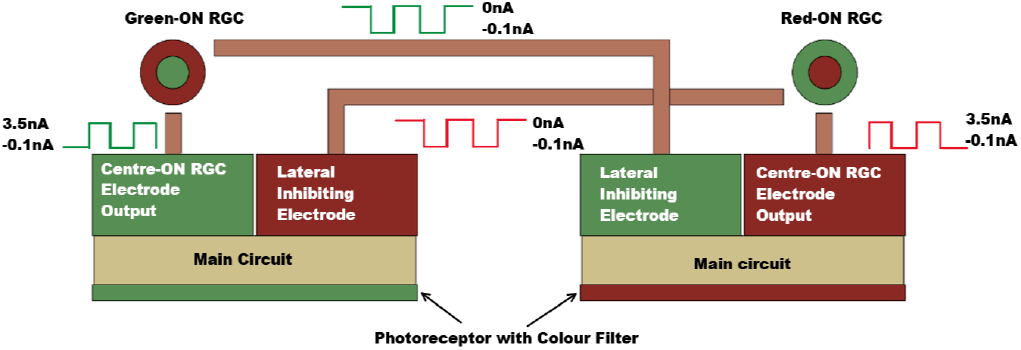
Lateral Inhibition via a Pixel pair replicating RGC opponent cell pairs. Each pixel has two electrodes (Fig. 6).

**Figure 8:**
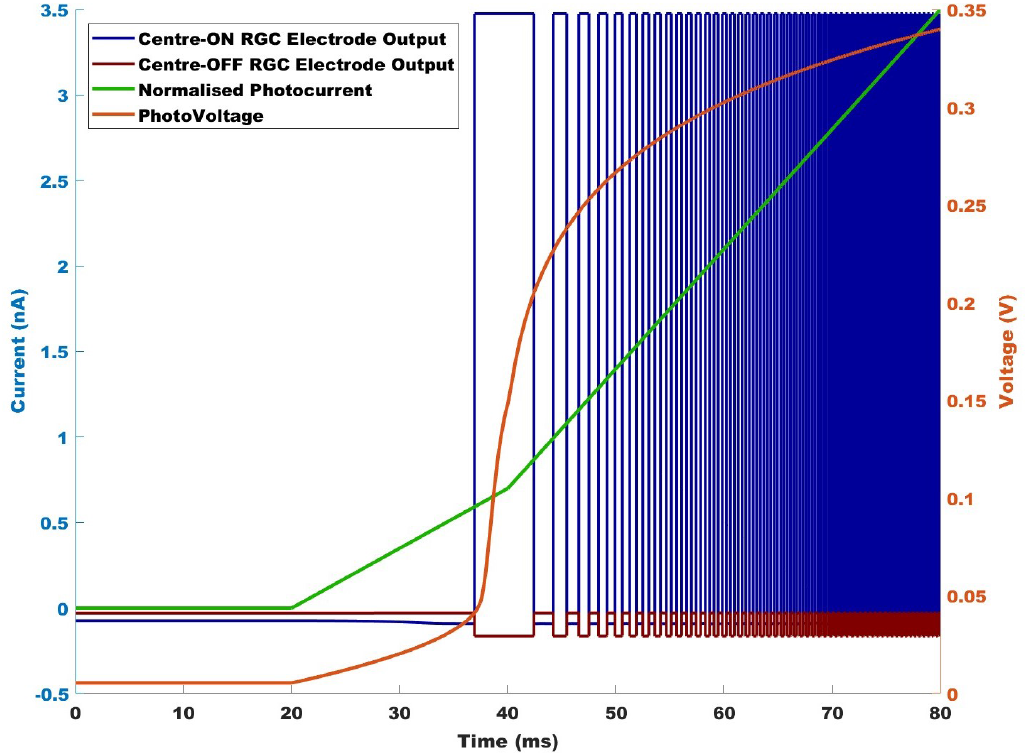
LTSpice Simulation Output at the Electrode. The photocurrent increases from 0nA to 100nA (resulting in a frequency increase) but has been normalised for illustration purposes.

## IV. Photo-receptive array development and method

The mapping of rods and cones in the natural (and artificial) retina is illustrated in Figure 9. Table II shows the properties of rods and cones within the retina, and compares them to both photodiodes and photo-transistors. Although both semiconductor devices are responsive to light, there is a key difference between them; a photodiode converts light energy into electrical energy, whereas a photo-transistor is a transistor that is sensitive to light and amplifies the current generated from light energy [20]. Photodiodes were selected to activate RGCs linked with cones, while phototransistors target RGCs associated with rods (Table II).

**Figure 9:**
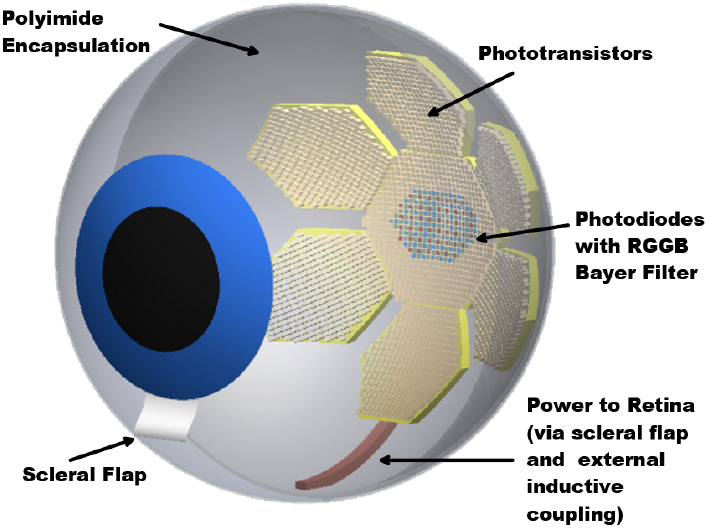
Not to scale CAD image of an eye fitted with an artificial curved retina (inspired by CurvIS [3]). The RGGB Bayer Patterned Coloured pixels (photodiodes) in the centre represents the fovea region with densely packed cone cells (responsible for colour and high acuity vision). The remainder of the pixels (photo-transistors) represent rod cells (responsible for low-light vision).

**Table II.**
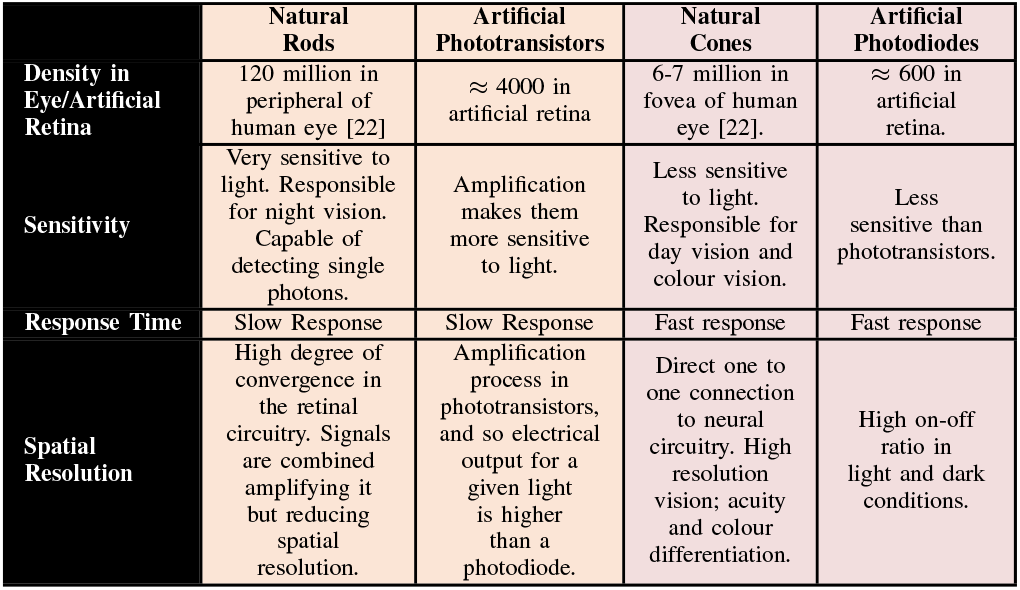
Comparison between natural and artificial photoreceptors [20].

### A. Photoreceptive Semiconductor Device Development

The P3HT:PCBM blend is widely used in organic photodiodes and phototransistors due to a combination of its electrical, optical, and physical properties. P3HT is a conjugated polymer that acts as a p-type semiconductor with good hole mobility. PCBM is a fullerene derivative that acts as an n-type semiconductor with good electron mobility. Together they form a bulk heterojunction (BHJ) structure (Fig. 10a), facilitating efficient electron-hole separation into free charge carriers upon light absorption [21]. The inclusion of interfacial layers such as PEDOT:PSS (anode) and Ca (cathode) in the photodiode facilitate transport and help align the energy levels between the electrodes and the active materials (Fig. 10b). Reducing the injection barriers results in a decrease of the threshold voltage. The choice of metal contacts has also been made to minimise the work function difference between the anode/cathode and the HUMO/LUMO levels of the active materials. Phototransistors require a high-quality dielectric to ensure a high capacitance per unit area, which can significantly lower the gate voltage required to turn on the device and induce the channel formation for charge transport. In this paper, 500nm of PMMA dielectric has been selected in order to increase capacitance (Fig. 11a).

**Figure 10:**
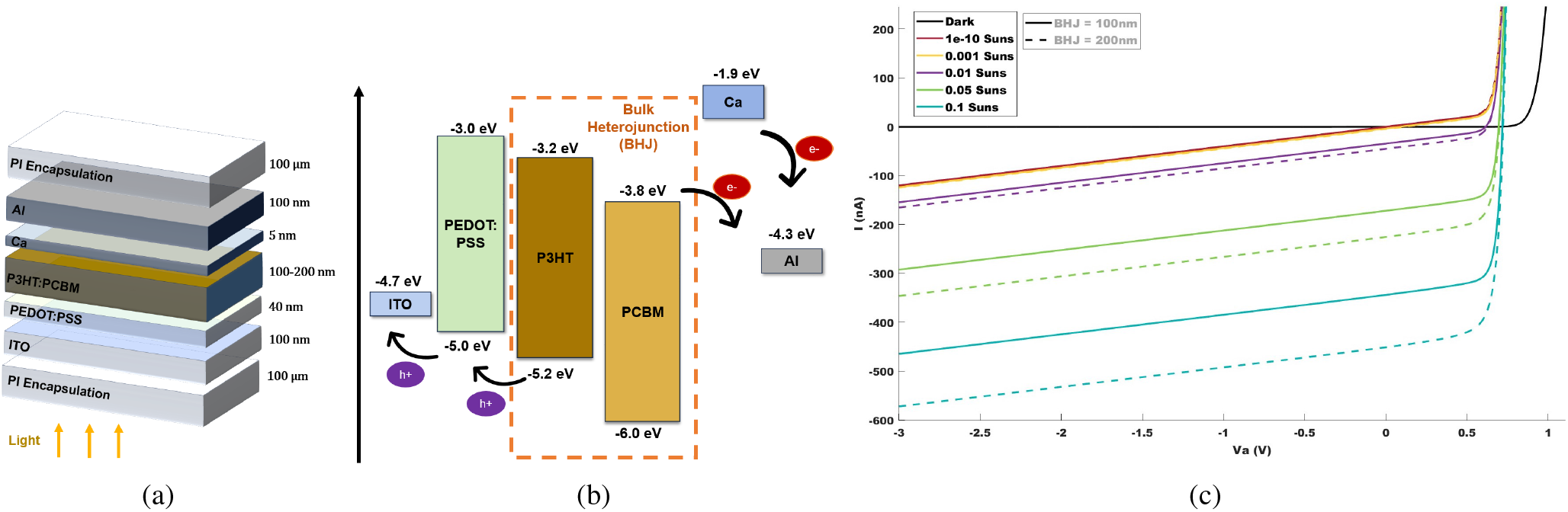
a) The semiconductor stack used in the photodiode simulations in OghmaNano. b) Energy Levels between contacts and active materials in the photodiode. Calcium is used as an electron transport layer, and PEDOT:PSS is used as a hole transport layer c) Output characteristics of a photodiode under increasing light intensity and different BHJ thicknesses. Photodiodes have a higher on/off ratio when operating in reverse bias. Note that 1*Suns* = 1000*W/m*^2^.

**Figure 11:**
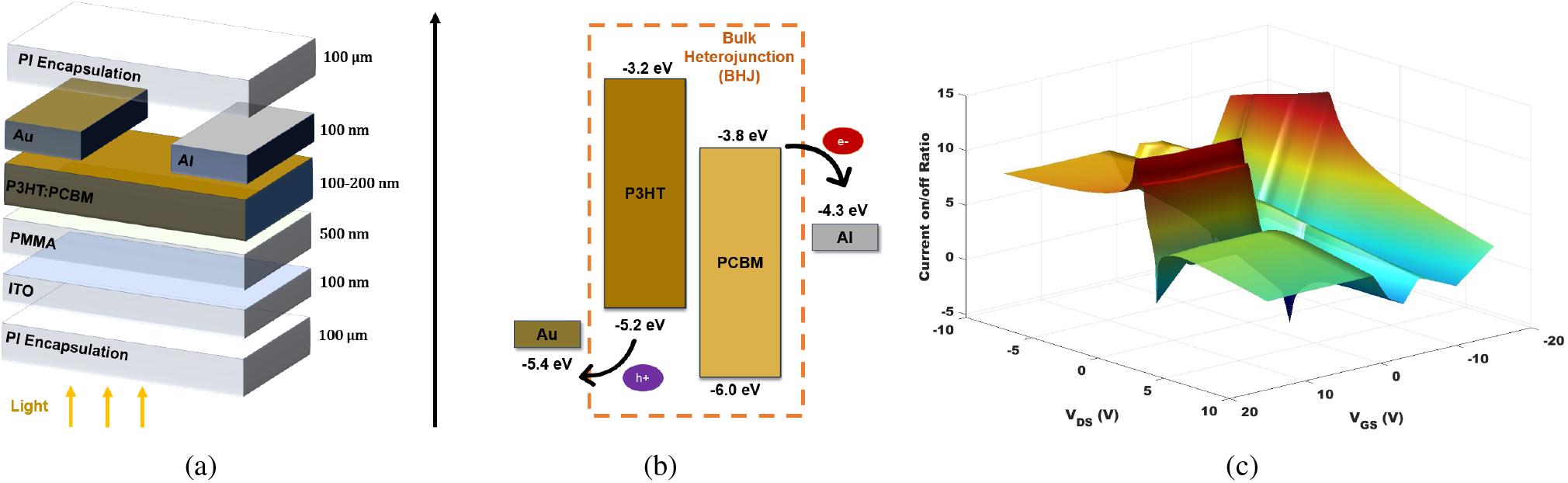
a) The semiconductor stack used in the phototransistor simulations in OghmaNano. ITO is used as the Gate in order to create the field effect and PMMA is the dielectric used. b) Energy Levels between contacts and active materials in the phototransistor. c) Surface plot to show a parameter sweep (*V*_*GS*_ and *V*_*DS*_), in order to optimise the on/off current ratio between Dark and 10^−3^*Suns*.

### B. Electrical Properties and Photo Response - OghmaNano Simulation Results

OghmaNano is a general purpose model for the simulation of opto-electronic devices [23] [24] [25]. The properties of both the photodiode and phototransistor were simulated in OghmaNano (Version 8.0.044) using finite element analysis (FEA) - see parameters in Table S4. The overall device geometry, including the thickness of the dielectric and semiconductor layers, is optimised to ensure that the electric field required to turn-on the device is minimised. For both devices, the substrate size in the simulations was set at 200*µm ×* 200*µm*. Considering the current density output of both devices, selecting a substrate size of 200*µm ×* 200*µm* yielded an optimal current output. Additionally, this size is compatible with the eye’s geometry and aligns with the optical analysis conducted by the developers of CurvIS [3].

Research shows that the minimum threshold light intensity for a human eye can be as little as ≈ 10^−19^*W/m*^2^, however a more commonly reported value is between 10^−12^ to 10^−10^*W/m*^2^ [26], which is equivalent to ≈ 10^−13^*Suns* where 1*Suns* = 1000*W/m*^2^.

The ability of the human eye to perceive very low levels of light is primarily attributed to the high sensitivity of rod cells. Similarly, the phototransistors developed for this study demonstrate a comparable response to minimal light intensities, as illustrated in Figures 12a and 12b. Compared to photodiodes, phototransistors can detect subtler variations in light intensity, although their response is logarithmic rather than linear, necessitating logarithmic interpretation of their output. This logarithmic scaling allows phototransistors to maintain sensitivity over a broader range of light intensities, with the scale of change increasing exponentially as light intensity rises. The phototransistor designed and simulated in this paper produces a very small current output of 1nA at ≈ 10^−4^*W/m*^2^ (≈ 10^−7^*Suns*). This light intensity is the first instance at which the current increases and a high enough on/off ratio exists between light and dark current output. Although higher than the absolute minimum that the human eye can detect, this is still an effective response for vision in the dark.

**Figure 12:**
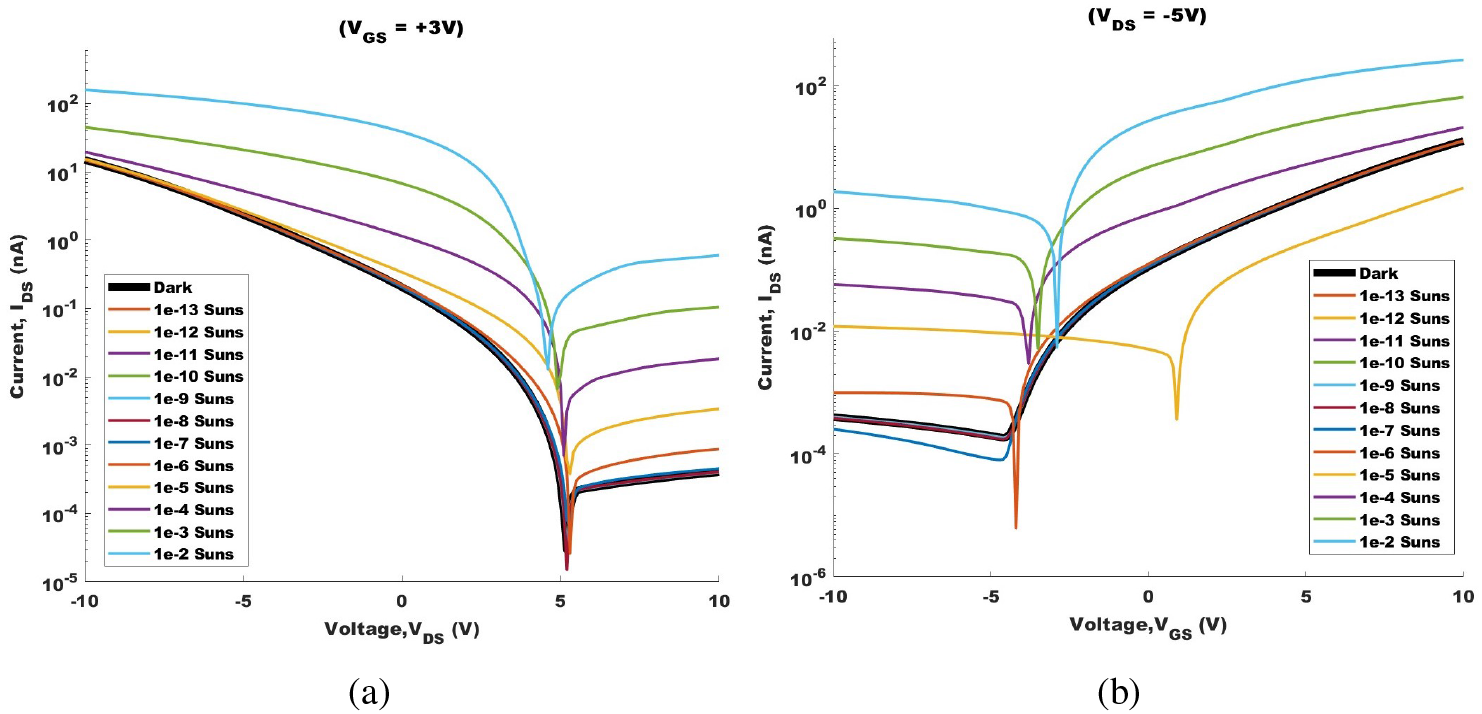
a) Output characteristics of a phototransistor under increasing light intensity at *V*_*GS*_ = 3*V*. b) Transfer characteristics of a phototransistor under increasing light intensity at *V*_*DS*_ = −5*V*. Note that 1*Suns* = 1000*W/m*^2^.

Owing to the BHJ blend of both p-type and n-type semiconductors, these phototransistors exhibit characteristics similar to ambipolar transistors. This enables them to operate effectively across both p-type and n-type regions, as shown in Figure 11c. The 3D surface plot illustrates the operating points that yield the highest and lowest on/off ratios under light and dark conditions. While operating at a positive drain-source voltage yields slightly higher output currents, it adversely affects the on/off ratio. Conversely, using a negative *V*_*DS*_ results in only a minimal reduction in current output while significantly improving the on/off ratio. Using a gate-source voltage (*V*_*GS*_) of ≈ -15V achieves the best on/off ratio, but such a high voltage may not be suitable for sensitive applications like an artificial retina. Instead, a more practical choice is a *V*_*GS*_ of 3V, which still offers a significantly high on/off ratio. By setting *V*_*GS*_ to 3V and *V*_*DS*_ to -5V, the phototransistor effectively operates in its saturation region as a p-type transistor.

The photodiodes, on the other hand, only initiate a response at ≈ 10^−10^*Suns* or ≈ 10^−7^*W/m*^2^. Figure 10c demonstrates a significant on/off ratio, where at an applied voltage of -3V (Reverse bias), the current shifts from 0nA (dark) to -100nA under very low illumination of ≈ 10^−10^*Suns*. Despite this substantial on/off ratio, the photodiode’s response remains consistent across low light intensities 10^−10^ to 10^−3^*Suns*. Subsequent changes are only visible at higher light intensities. Although this range is not as low as the minimum light intensity perceivable by the human eye (energy equivalent to a single photon [26]), it proves adequate for cone cells. Cone cells primarily function in daylight and colour vision. Indoor lighting typically ranges from 1*W/m*^2^ (0.001*Suns*) to 100*W/m*^2^ (0.1*Suns*) depending on the light sources. Figure 10c shows that photodiodes are the optimal choice for cone cells due to their ability to exhibit significant current shifts across various light intensities in this range. In contrast to phototransistors, photodiodes typically exhibit a more linear change (Fig. 10c), where the response increases proportionally with the light intensity. For the best performance, the photodiodes should be operated in reverse bias for the highest on/off ratio with an applied voltage between -2V to -3V. A forward biased diode does not have much significance in terms of its photo-response.

## V. Proposed Solution of an Intraocular Artificial Retina

### A. LTSpice Circuit Simulations and Optimisation

Research shows that the responsiveness of RGCs varies across cell types with a range of stimulation frequencies from 20Hz to 700Hz [11]. Assuming this range is correct for most RGCs, the proposed circuit shows an increase in stimulation frequency with photocurrent.

Figure 13a indicates that a capacitance value of 1nF for C2 yields an output frequency range that most suitably corresponds to the response range of RGCs in the human eye as well as the photodiode output characteristics. Additionally, optimisation of the current sink’s aspect ratio, (*W/L*)_*M*5_, substantially improves the on/off ratio, particularly at lower currents. It is observed that a smaller aspect ratio of M5 leads to a higher frequency output for photocurrents below 150nA, enhancing the circuit’s responsiveness and sensitivity. Post optimisation, the circuit produces a frequency output of 2.5Hz at 0nA (within background baseline frequencies) and 30Hz at 50nA. At 100nA, to which the photodiode responds to 1*W/m*^2^ (0.001*Suns*) light intensity, the output frequency is ≈ 60Hz. At 600nA, the frequency rises to 600Hz in response to 100*W/m*^2^ (0.1*Suns*).

**Figure 13:**
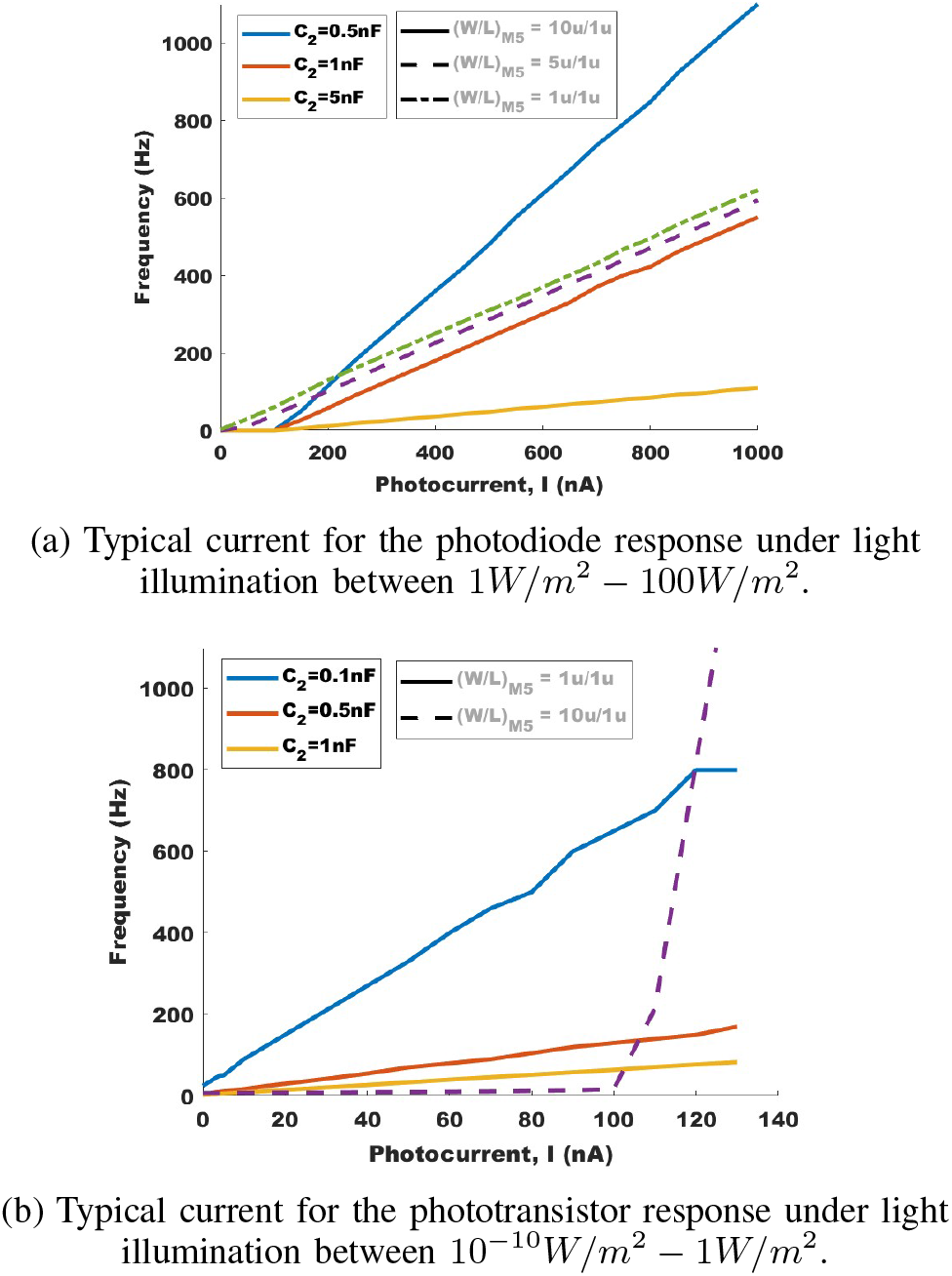
Circuit Output frequency against photocurrent input, for parameter optimisation. M5 is the current sink that regulates the photo-voltage. C2 determines the pulse train frequency.

For the phototransistor however, a capacitance value of 0.1nF (Figure 13b) yields an output frequency range that aligns well with the sensitivity of rods and the current output characteristics of the phototransistor itself. As demonstrated in the phototransistor’s output and transfer characteristics (Fig. 12a and 12b), there exists a non-linear relationship between current increase and light intensity. To enhance amplification of the photocurrent and subsequent photo-voltage output, positive feedback is introduced between the output of the transimpedance amplifier (Drain of M5) and the input (Gate of M1). To accommodate the wide range of non-linear photocurrents, the circuit is optimised to deliver a linear frequency output. This ensures that even very small currents, such as 1nA at ≈ 10^−4^*W/m*^2^, can be effectively amplified to produce a frequency output of 25Hz, replicating the high sensitivity of rod cells. However, as light intensity increases beyond a certain threshold, e.g., at 0.1*W/m*^2^, the frequency output starts to plateau, reaching a maximum of 700Hz. This reflects the reduced effectiveness of rods under brighter light conditions.

### B. Implementation

Powering the electronic circuitry within the eye presents a challenge. The Alpha-IMS subretinal implant [2] addressed this by using a coil and inductive power transmission housed in a ceramic casing positioned subdermally behind the ear (Fig. S5). This system can potentially be designed as a portable handheld battery-powered device with an electromagnetic coil similar to the Alpha IMS. The external coil is magnetically secured in place to enable “wireless” power transmission to the implant through the skull. The internal component of the inductive coupling device contains an electronic power-converting circuit to meet the circuit’s multiple voltage supply requirements. Power wires will then travel from the inductive coupling area to the artificial retina within the eye, via the scleral flap (Fig. 9).

## VI. Discussion

### A. Biological Limitations

For this bio-electronic project to reach its full potential, a deeper understanding of RGC neuronal signal spikes is essential. Currently, the precise relationship between the frequency of these spikes and varying light intensities remains undefined. In practice, RGCs exhibit a diverse range of responses to light: some prioritise capturing fine details through slower, more prolonged signalling, while others rapidly adjust their firing rates to track significant shifts in brightness and motion. Simulation studies, such as those conducted with NEURON in this paper, indicate that neuron action potentials maintain a consistent shape, with variations primarily in spike frequency. However, in the proposed circuit, the pulse width itself changes in order to adjust the output frequency. At this stage, the impact of this is unclear; some studies suggest that varying the pulse width could potentially impact cell thresholds and signal amplitude [11]. An aspect of neuronal behaviour not explored in this study is the accommodation property. Accommodation in the context of neuron physiology refers to a neuron’s ability to adjust its responsiveness over time [18] to a constant or repetitive stimulus. This can include changes in the neuron’s threshold for firing action potentials in response to sustained inputs. Stimulating RGCs in a precise and controlled manner often demands very small currents, particularly in a direct and highly localised manner. NEURON simulations demonstrate that the current required to trigger an action potential can indeed be in the nanoampere (nA) range, depending on various factors including cell-specific biophysics and morphology. These parameters influence the threshold requirements, emphasising the need for specific parameters tailored to each RGC type within the retina [11].

### B. Future Work

Further development will focus on optimising semiconductor layer thicknesses within the photodiodes and phototransistors, and investigating alternative active materials such as *MoS*_2_ and Graphene as proposed by CurvIS [3]. However, these new propositions present challenges in simulation and manufacturing due to their novelty. In this study, the widely recognised photoreceptive semiconductor BHJ blend of P3HT:PCBM was selected to demonstrate the project’s feasibility and reduce the risks associated with the exploration of new materials. OghmaNano is primarily used for solar cell development and the material database for P3HT:PCBM is detailed. Since CurvIS and other studies have demonstrated feasible semiconductor manufacturing on curved substrates, further OghmaNano simulations may include FEA on a curved surface to account for any geometry-dependent changes in photoreceptor output.

Simulating the circuit FETs using biocompatible electronics parameters, rather than ideal LTSpice models, is essential to validate the design for fabrication. Future iterations may also incorporate feedback mechanisms to dynamically adjust the electrode output frequency, accommodating human variability and enabling restoration of RGC resting potential through optimised biphasic pulsing.

## VII. Conclusion

This project aimed to create a comprehensive solution for stimulating neuronal signals in RGCs using a curved array and a circuit that produces an output backed by NEURON simulations. While CurvIS has developed phototransistors, further work was required to optimise and miniaturise the signal processing components. The circuit design proposed in this paper addresses miniaturisation challenges by eliminating the need for separate microcontrollers for each photoreceptive pixel, as proposed by CurvIS. In summary, this project represents a significant advancement towards constructing an artificial retina for individuals with visual impairments. NEURON simulations have established signal thresholds for RGCs, providing a baseline for future research. The proposed circuit offers flexibility to adjust parameters for accurate signal transmission, while also integrating innovative strategies to mitigate cross-talk and facilitate physical implementation into a retina. Although there is still a requirement for additional refinement in the photoreceptor semiconductor chemistry, frequency output, and geometry optimisation, the proposed solution in this project should be considered for further development.

## Supporting information

Main Manuscript Only

Supplementary Info Only

## Supplementary Information

### I. NEURON Simulations

GitHub Link to the NEURON (Version 8.2) Code used in the simulations in this paper: https://github.com/VedikaBedi/NEURON_Code_VedikaBedi/tree/main

**Table S1.**
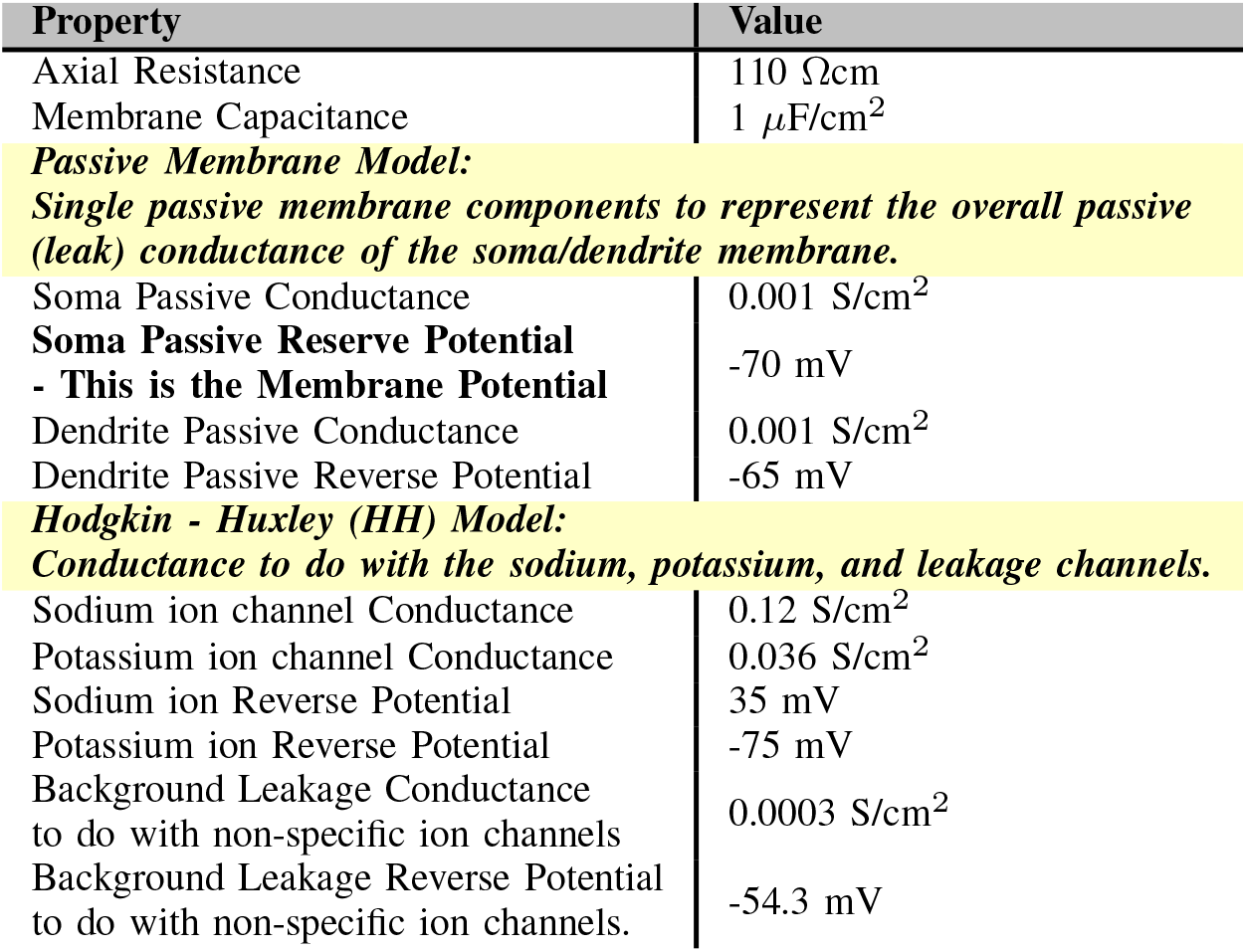
Biophysical parameters used in NEURON 8.2 [1]. In NEURON, the RGC cell was modelled with; soma (D = 5 *µm*), a dendrite synapsing with a bipolar cell (D = 0.3 *µm*, L = 17 *µm*M), and the axon split into three parts consisting of the axon hillock (AH: D = 1.25 *µm*, L = 40 *µm*), Axon Initial segment, (AIS: D = 0.60 *µm*, L = 40 *µm*), and the remainder of the Axon (D=0.91*µm*, L= 1170 *µm*) [1].

**Figure S1:**
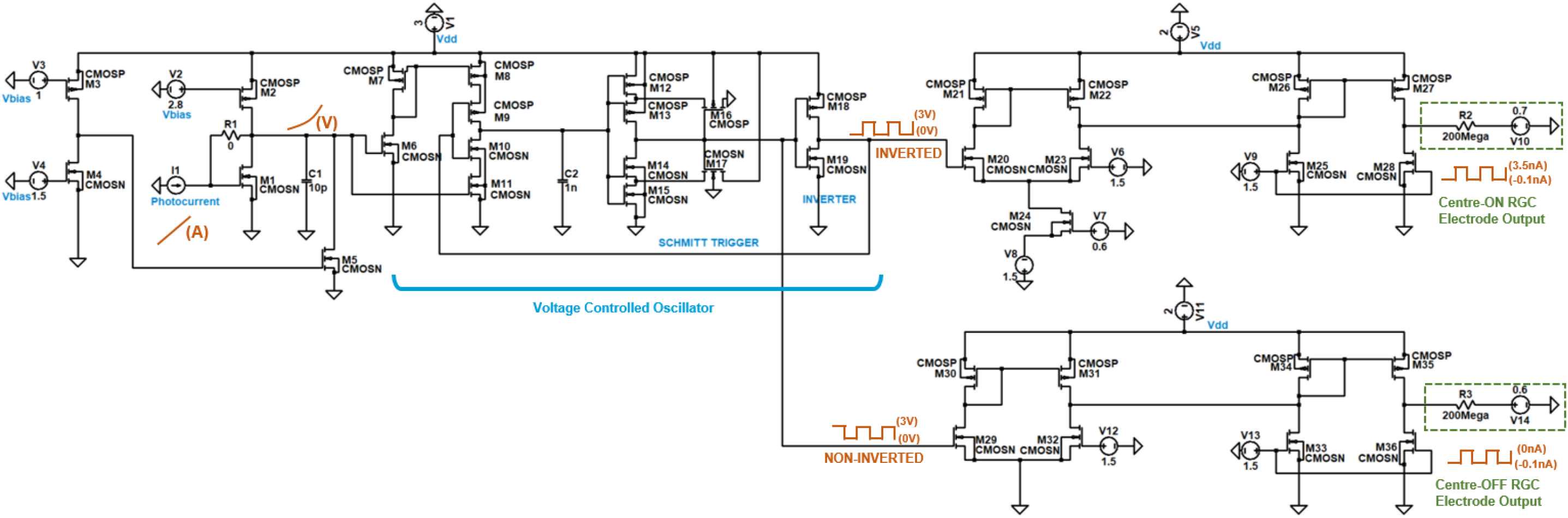
Proposed Circuit (enlarged figure)

### II. Circuit Modelling and LTSpice Simulations

**Table S2.**
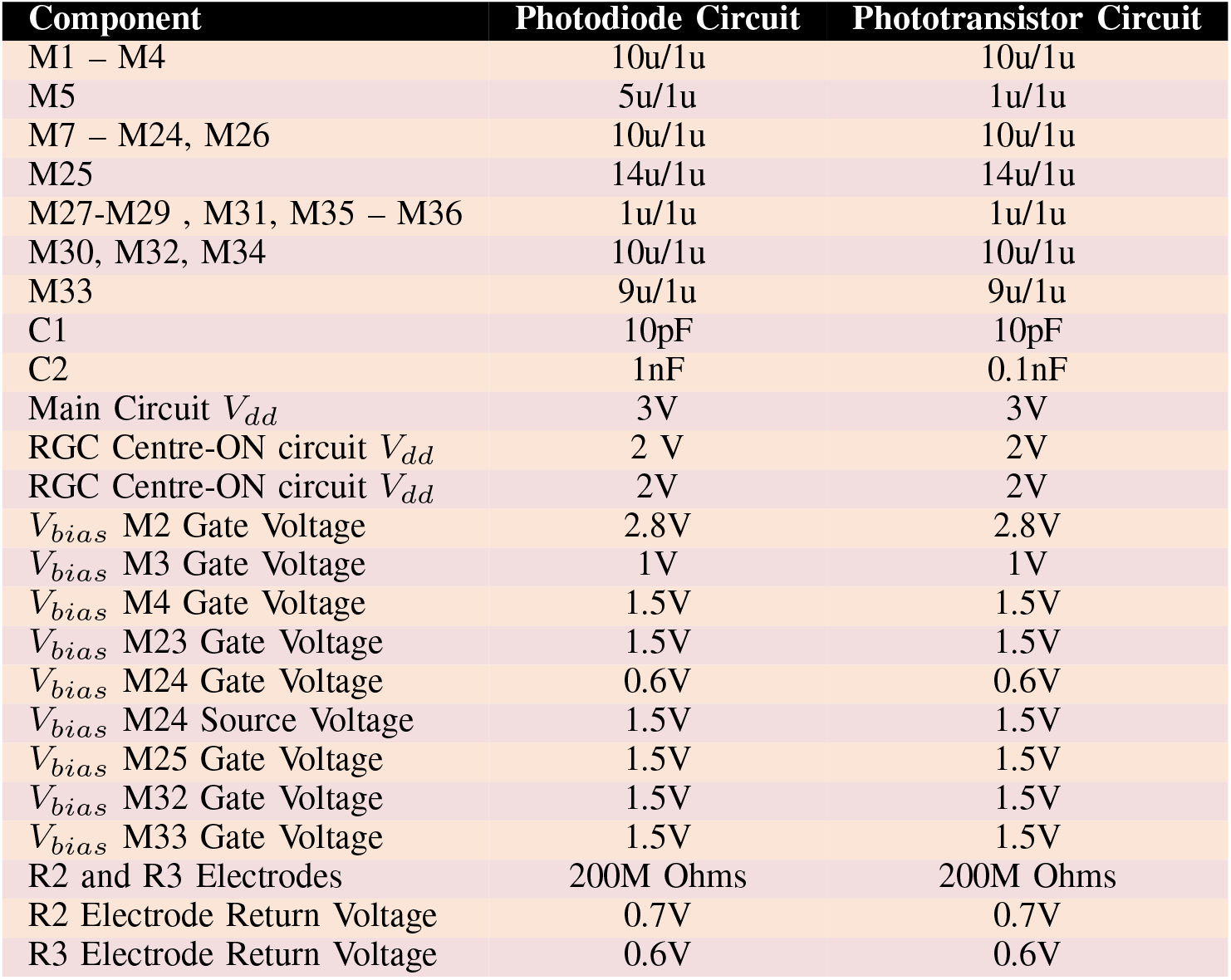
Breakdown of components values in the proposed neuron spiking circuit that converts a photocurrent to a suitable output current meeting the threshold requirements for RGCs.

**Table S3.**
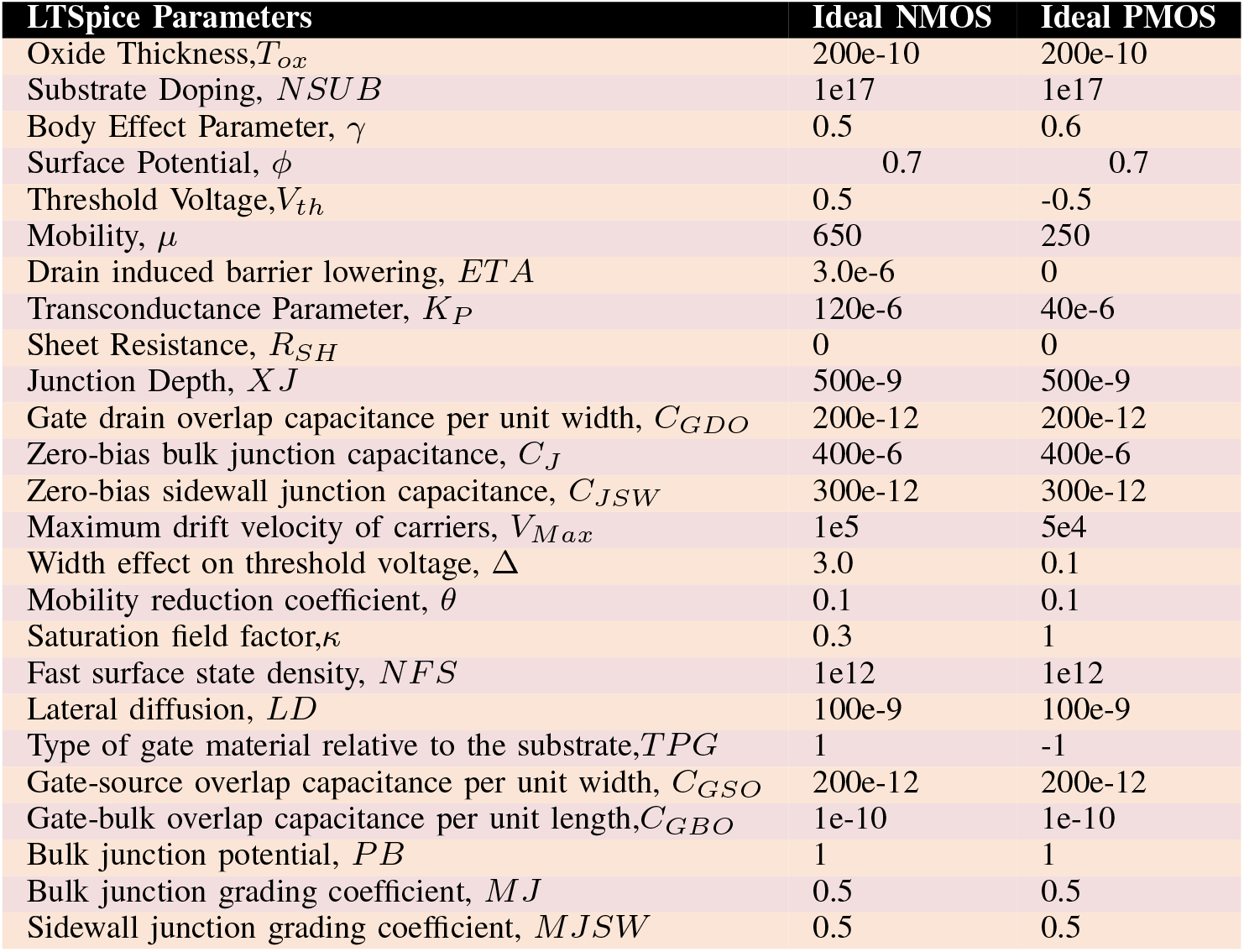
MOSFET parameters used in the Level 3 model in LTSpice (Version 24.0.9).

**Figure S2:**
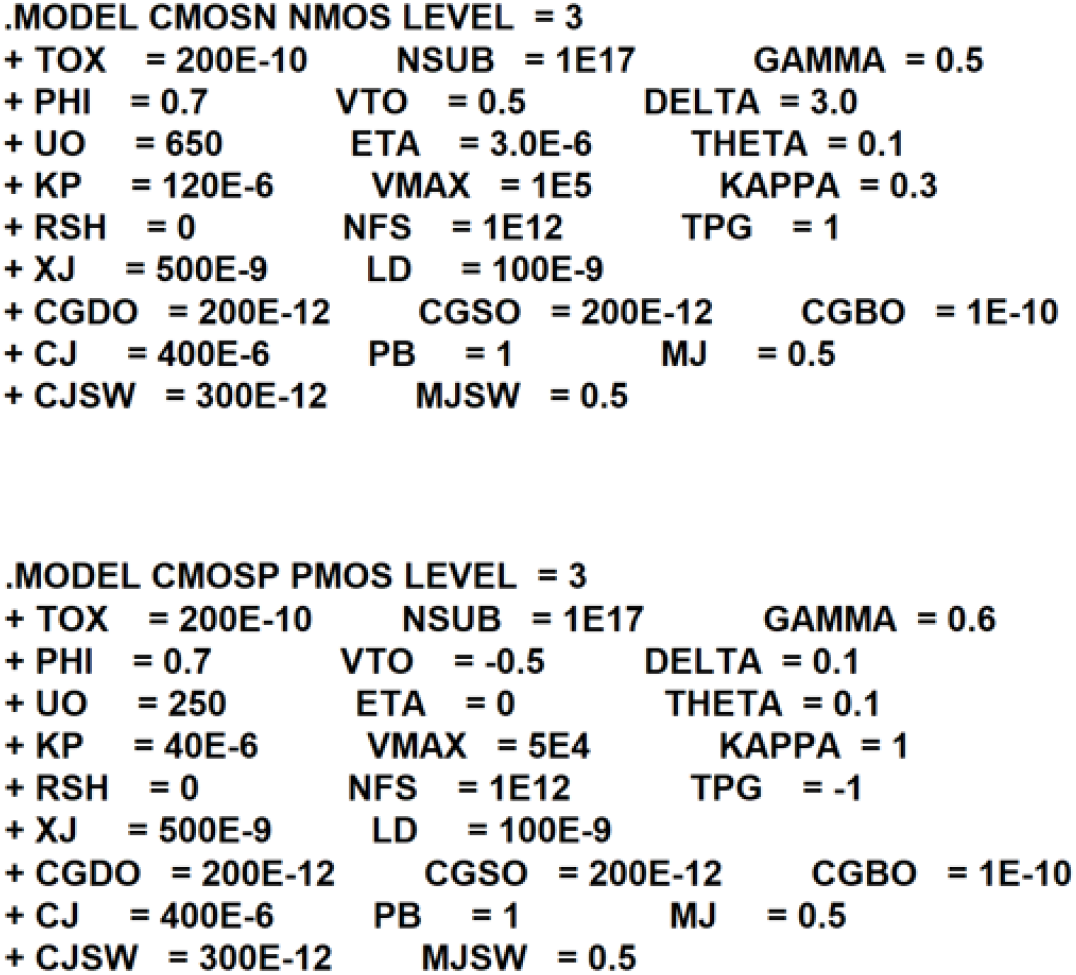
.MODEL statements in LTSpice to set up the Level 3 MOSFET model

### III. OghmaNano Simulations

**Table S4.**
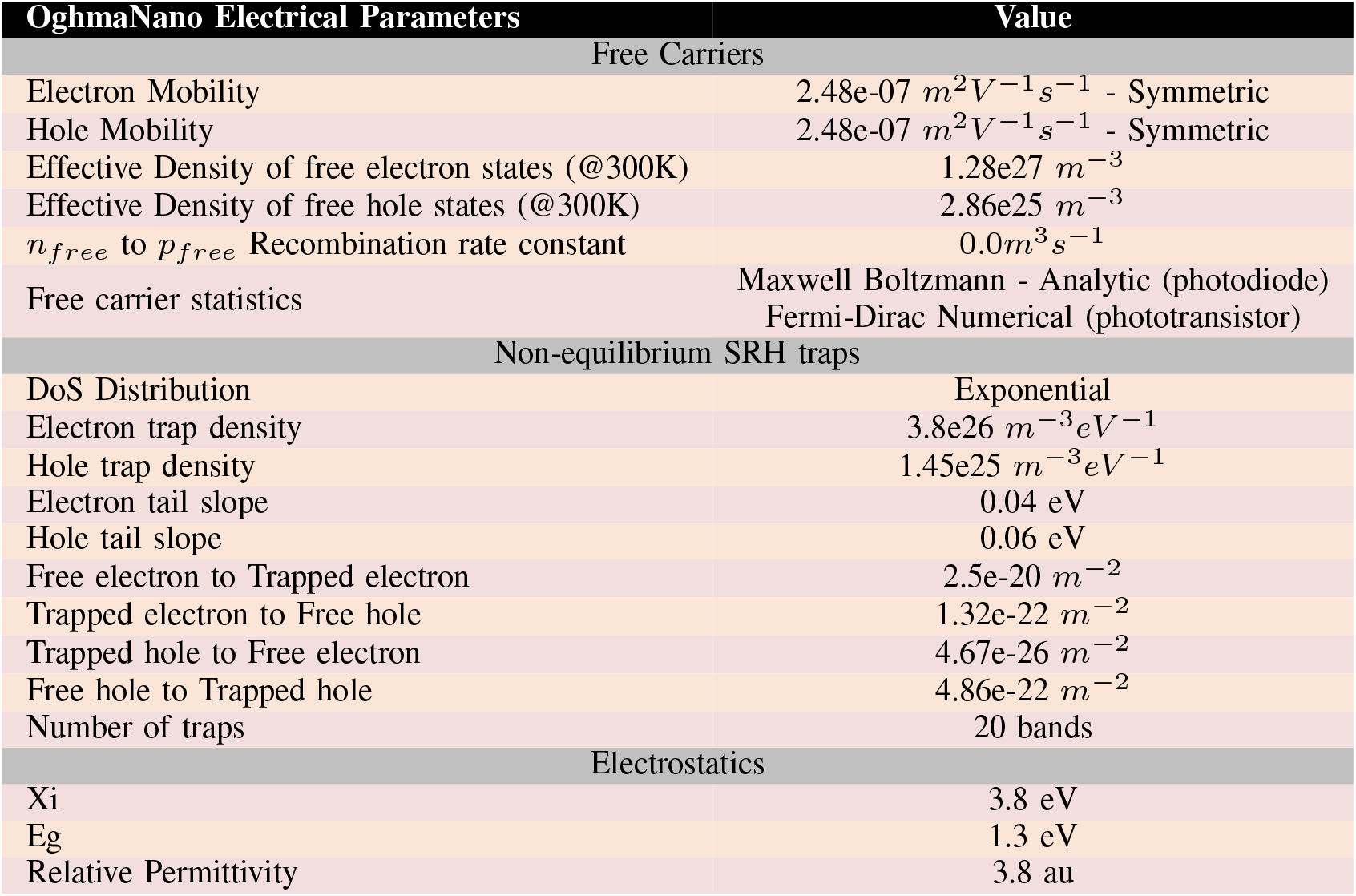
OghmaNano (Version 8.0.044) Simulation Parameters - Electrical Parameters of the active material (P3HT:PCBM)

| OghmaNano Electrical Parameters | Value |
| --- | --- |
| Free Carriers |  |
| Electron Mobility | 2.48e-07 $m^2V^{-1}s^{-1}$ - Symmetric |
| Hole Mobility | 2.48e-07 $m^2V^{-1}s^{-1}$ - Symmetric |
| Effective Density of free electron states (@300K) | 1.28e27 $m^{-3}$ |
| Effective Density of free hole states (@300K) | 2.86e25 $m^{-3}$ |
| $n_{free}$ to $p_{free}$ Recombination rate constant | 0.0 $m^3s^{-1}$ |
| Free carrier statistics | Maxwell Boltzmann - Analytic (photodiode)<br>Fermi-Dirac Numerical (phototransistor) |
| Non-equilibrium SRH traps |  |
| DoS Distribution | Exponential |
| Electron trap density | 3.8e26 $m^{-3}eV^{-1}$ |
| Hole trap density | 1.45e25 $m^{-3}eV^{-1}$ |
| Electron tail slope | 0.04 eV |
| Hole tail slope | 0.06 eV |
| Free electron to Trapped electron | 2.5e-20 $m^{-2}$ |
| Trapped electron to Free hole | 1.32e-22 $m^{-2}$ |
| Trapped hole to Free electron | 4.67e-26 $m^{-2}$ |
| Free hole to Trapped hole | 4.86e-22 $m^{-2}$ |
| Number of traps | 20 bands |
| Electrostatics |  |
| Xi | 3.8 eV |
| Eg | 1.3 eV |
| Relative Permittivity | 3.8 au |

**Figure S3:**
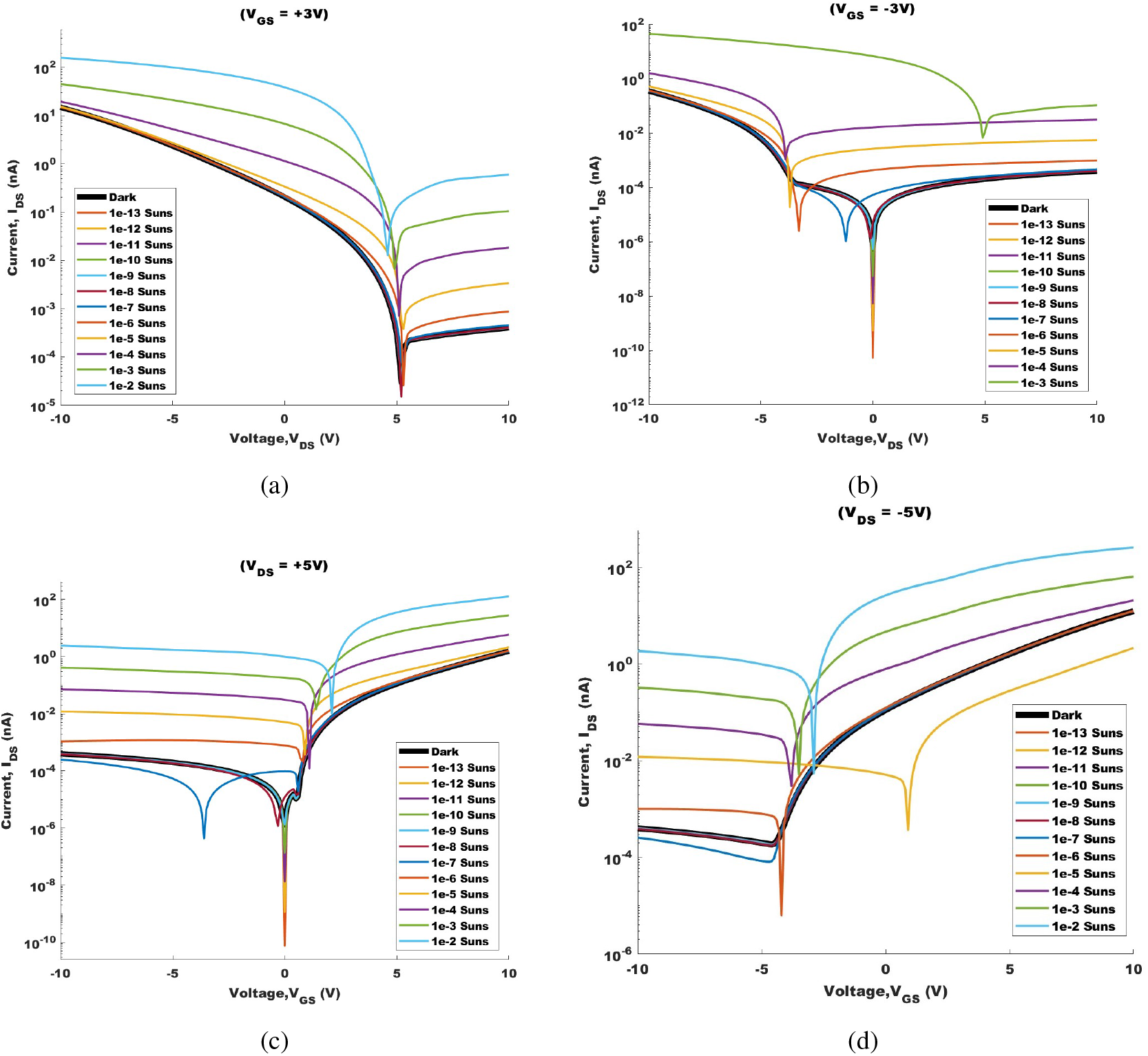
Output and Transfer Characteristics of the phototransistor. This provides better illustration of how the choice of positive and negative voltages impacts the characteristics of the phototransistor. Note this is due to an unequal blend of P3HT:PCBM being present in the semiconductor. This material database has been taken from a solar cell model, and so the concentration blend is yet to be optimised.

**Figure S4:**
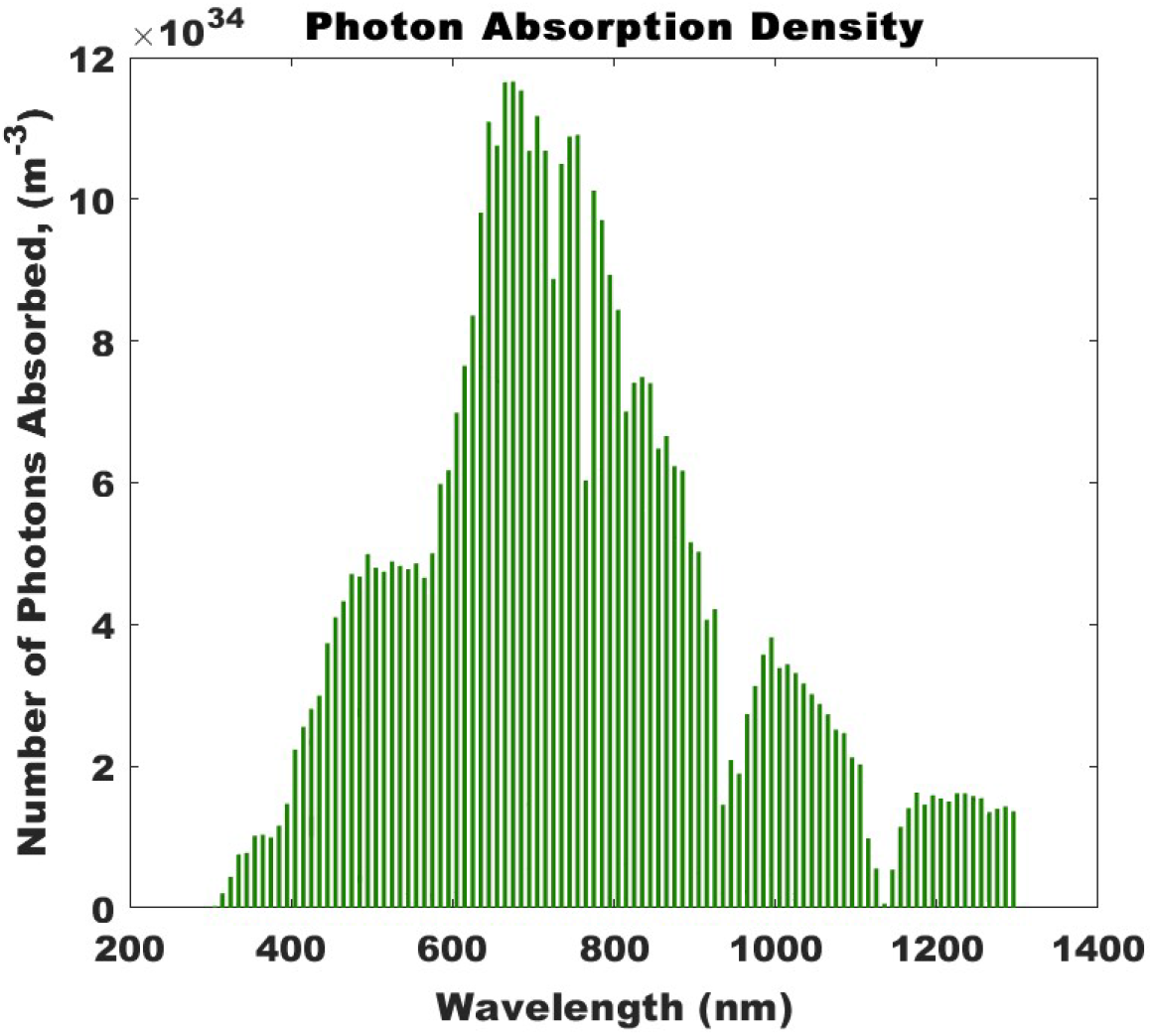
The P3HT:PCBM blend has excellent photon absorptivity at all visible light wavelengths, with the highest absorption at 650nm. The photoreceptors will however, require an Infrared filter as it also is shown to have good absorption between 700-1200nm.

**Figure S5:**
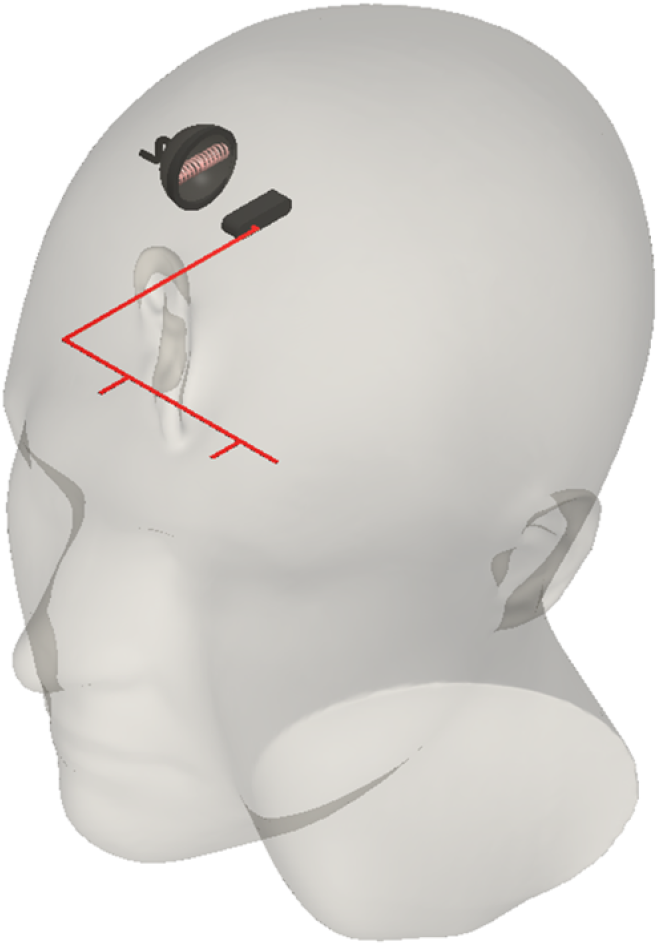
Not to Scale CAD image of a human head fitted with an inductive coupling device with power wires for each eye.

