## Supplementary material for "An Integrated Photoreceptor-to-RGC Stimulation Circuit for Intraocular Visual Prostheses": Main Manuscript Only

Vedika Bedi supervised by Dr. Mujeeb U. Chaudhry

*Abstract*—**Visual prostheses face a critical miniaturisation challenge: converting photoreceptor signals to biologically appropriate retinal ganglion cell (RGC) stimulation patterns within the spatial constraints of intraocular implants. Existing systems rely on external microcontrollers for signal processing, limiting scalability for high-density pixel arrays. This paper presents an integrated per-pixel circuit architecture that directly converts photocurrent into frequency-modulated current pulses that match RGC activation thresholds.**

**The design targets are established through NEURON computational modelling of red-green colour-opponent midget RGCs, identifying stimulation thresholds of +0.1nA to +3.5nA for depolarisation and -0.1nA for repolarisation. The proposed circuit combines a transimpedance amplifier, a voltage-controlled oscillator with a Schmitt trigger, and a current-controlled output stage to generate biphasic pulses within these thresholds. A complementary output provides lateral inhibition, reducing crosstalk between adjacent RGC stimulation sites.**

**Photoreceptor integration is achieved using P3HT:PCBM organic photodiodes for cone-associated RGCs and phototransistors for rod-associated RGCs, validated through OghmaNano finite element simulations. The photodiode circuit produces output frequencies of 2.5Hz (dark) to 600Hz (100 $W/m^2$), matching reported RGC response ranges. This architecture eliminates external processing requirements, enabling scalable high-density retinal prostheses design.**

*Index Terms*—**Retinal Prosthesis, Retinal Ganglion Cells, Neural Stimulation Circuits, Organic Photodetectors, Visual Prosthesis, Neuromorphic Circuits**

*The authors would like to thank EPSRC (New Investigator Award EP/V037862/1) and the TUBITAK 2221 programme for the financial support.* *Vedika Bedi was with Durham University, Durham, DH1 3LE UK. She is now with University College London, London, WC1E 6BT UK. Dr. Mujeeb U. Chaudhry was with Durham University, Durham, DH1 3LE UK. He is now with Konya Technical University, Konya, Turkey.*

Figure 4 reveals that increasing pulse duration produces an apparent "saturation" effect, attributable to excessive depolarisation. A 1-2ms current pulse shows a characteristic spike followed by rapid decline, indicating the HH-modelled potassium ion release for repolarisation. However, this natural repolarisation mechanism proves insufficient for longer current pulses, establishing the need for an active inhibiting signal in the artificial stimulation circuit.

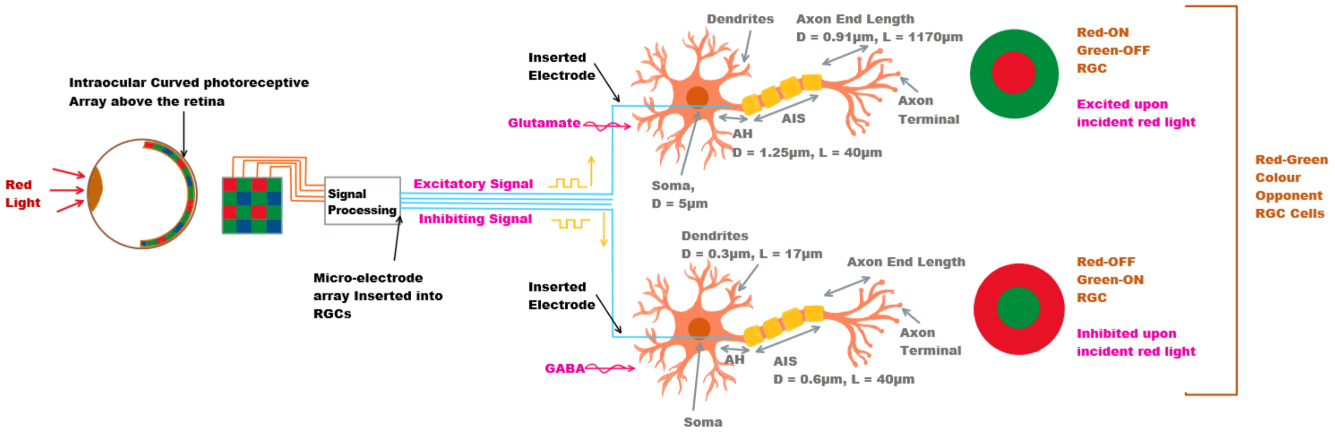


Figure 1: An illustration of the intraocular photoreceptive array inducing both excitatory and inhibitory signals in red-green colour opponent cells upon incident red light. For both, electrodes are shown to be inserted into the RGC cell axon and stimulated using a pulse via a function generator. Glutamate and GABA are illustrated to show synaptic neurotransmission under normal retinal operation. The neuron model shows the morphology of the RGC; where the soma is the cell body containing the nucleus, the dendrites receive inputs from other cells, and the axon carries the electrical impulse to be received by other neurons. [8].

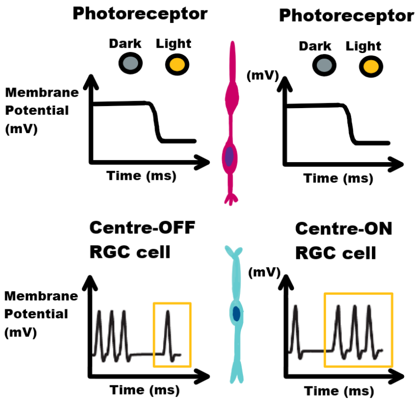


Figure 2: All photoreceptors respond with hyperpolarisation when transitioning from dark to illuminated conditions. Centre-OFF RGCs elevate their firing rates in darkness and diminish them in response to light stimuli. Conversely, Centre-ON RGCs increase their firing rates upon exposure to light and decrease them in darkness. [14]

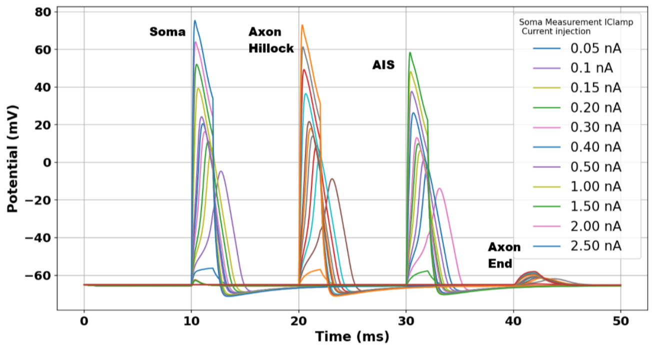


Figure 3: This graph from NEURON simulations shows the potential at the Soma, Axon Hillock, AIS, and Axon End, each with a varying current injection from 0.05 nA to 2.50 nA

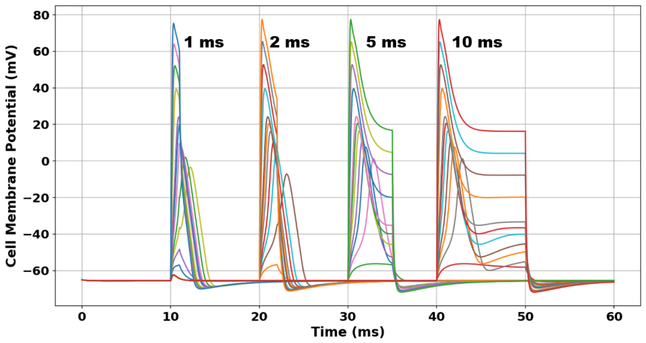


Figure 4: Graph from NEURON simulations shows the soma membrane potential and current injection but with varying levels of pulse duration (1ms, 2ms, 5ms, 10ms).

Biphasic stimulation pulses are therefore essential (Fig. 5). The cathodic phase depolarises the membrane and elicits a neural response, while a subsequent anodic phase discharges the accumulated charge around the electrode. Although some studies employ an equal anodic and cathodic current injection, NEURON results and proposed neuron circuit models such as the Izhikevich model [18], suggest a slow repolarisation aiming to prevent intense hyperpolarisation.

Table I summarises the membrane potential thresholds and the equivalent current pulses, establishing the design parameters for the proposed circuit. Depolarisation begins to occur at a threshold of approximately -55mV (Fig. 3) [8]. The neuron completely depolarises at a membrane potential of +80mV once the sodium channels are fully open, requiring a cathodic pulse of 2.5nA when artificially stimulated. For repolarisation, however, an anodic pulse of -0.1nA is sufficient (Fig. 5).

These four parameters — minimum depolarisation cur-

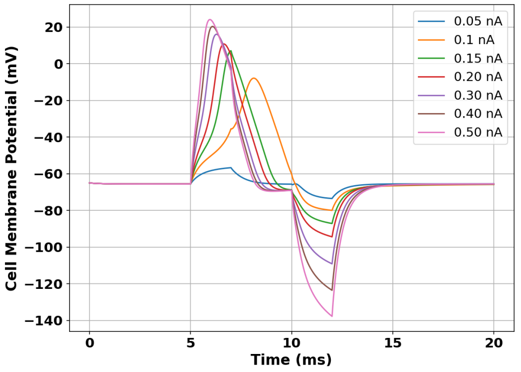


Figure 5: NEURON results from a biphasic pulse current injection. The first 2ms phase of the pulse (cathodic current) depolarises the cell membrane and elicits a natural response. The second 2ms phase (anodic current) balances the first phase to prevent net charge accumulating on the electrode. An intraphase delay of 2ms separates the pulses so that no reversal of the physiological effects occur [19].

rent (+0.1nA), maximum depolarisation current (+2.5nA), repolarisation current (-0.1nA), and frequency range (20-700Hz) — form the specifications for the stimulation circuit presented in Section III.

Table I
CURRENT PULSE REQUIREMENTS

| Phase | Membrane Potential | Current | Purpose |
|---|---|---|---|
| Resting Potential (Table S1) | -70mV | 0nA | Natural State (no visual stimuli) |
| Depolarisation Threshold [8] | -55mV (Fig. 3) | + 0.1nA | Minimum to induce ionic response [8] |
| Fully Depolarised Neuron | +40mV to +80mV (Fig. 3) | + 3.5nA | Maximum current before hyperpolarisation |
| Repolarisation (Fig. 5) | -70mV | - 0.1nA | Restores potential / Lateral Inhibition |

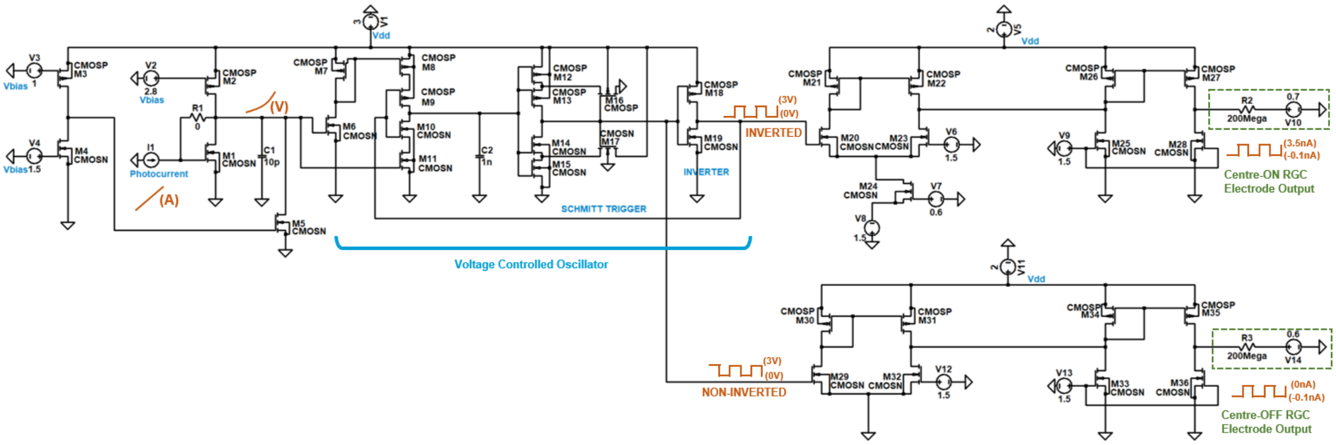


Figure 6: Proposed neuron spiking circuit (enlarged in Fig. S1) that converts a photocurrent to a suitable output current meeting the threshold requirements for RGCs. The electrodes (R2 and R3) induce an electrochemical response in the RGCs upon 'firing'.

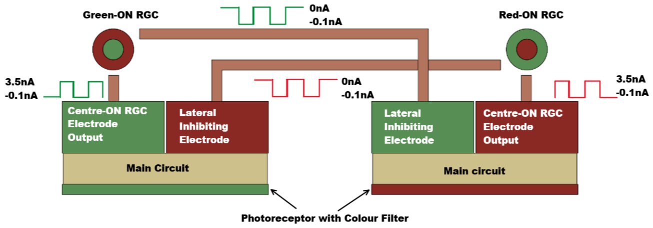


Figure 7: Lateral Inhibition via a Pixel pair replicating RGC opponent cell pairs. Each pixel has two electrodes (Fig. 6).

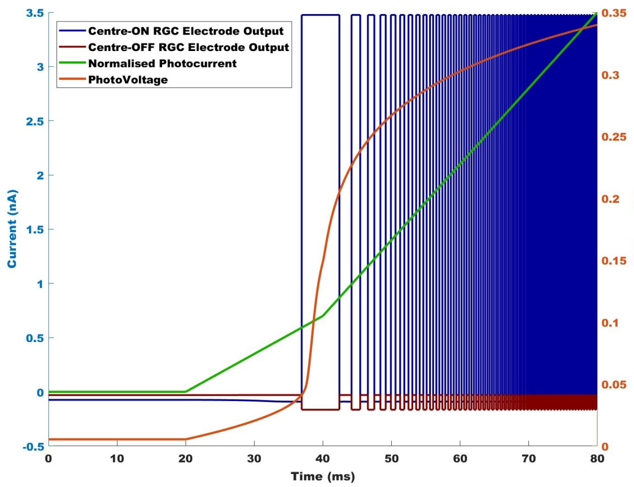


Figure 8: LTSpice Simulation Output at the Electrode. The photocurrent increases from 0nA to 100nA (resulting in a frequency increase) but has been normalised for illustration purposes.

is then output at a high level (0A). This lateral inhibition system operates as seen in Figure 7 and utilises pixel pairs based on RGC opponent cell pairs.

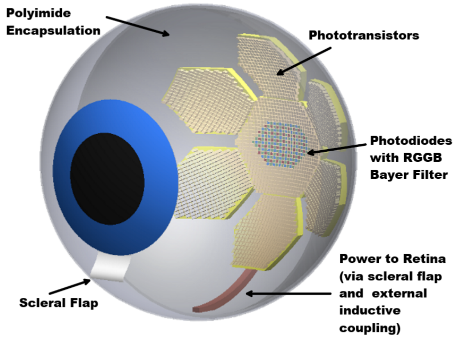


Figure 9: Not to scale CAD image of an eye fitted with an artificial curved retina (inspired by CurvIS [3]). The RGGB Bayer Patterned Coloured pixels (photodiodes) in the centre represents the fovea region with densely packed cone cells (responsible for colour and high acuity vision). The remainder of the pixels (photo-transistors) represent rod cells (responsible for low-light vision).

### IV. Photo-receptive Array Development and Method

The mapping of rods and cones in the natural (and artificial) retina is illustrated in Figure 9. Table II shows the properties of rods and cones within the retina, and compares them to both photodiodes and photo-transistors. Although both semiconductor devices are responsive to light, there is a key difference between them; a photodiode converts light energy into electrical energy, whereas a photo-transistor is a transistor that is sensitive to light and amplifies the current generated from light energy [20]. Photodiodes were selected to activate RGCs linked with cones, while phototransistors target RGCs associated with rods (Table II).

#### A. *Photoreceptive Semiconductor Device Development*

The P3HT:PCBM blend is widely used in organic photodiodes and phototransistors due to a combination of its electrical, optical, and physical properties. P3HT is a conjugated polymer that acts as a p-type semiconductor with good hole mobility. PCBM is a fullerene derivative that acts as an

Table II
COMPARISON BETWEEN NATURAL AND ARTIFICIAL PHOTORECEPTORS [20].

| | Natural Rods | Artificial Phototransistors | Natural Cones | Artificial Photodiodes |
|---|---|---|---|---|
| **Density in Eye/Artificial Retina** | 120 million in peripheral of human eye [22] | ≈ 4000 in artificial retina | 6-7 million in fovea of human eye [22]. | ≈ 600 in artificial retina. |
| **Sensitivity** | Very sensitive to light. Responsible for night vision. Capable of detecting single photons. | Amplification makes them more sensitive to light. | Less sensitive to light. Responsible for day vision and colour vision. | Less sensitive than phototransistors. |
| **Response Time** | Slow Response | Slow Response | Fast response | Fast response |
| **Spatial Resolution** | High degree of convergence in the retinal circuitry. Signals are combined amplifying it but reducing spatial resolution. | Amplification process in phototransistors, and so electrical output for a given light is higher than a photodiode. | Direct one to one connection to neural circuitry. High resolution vision; acuity and colour differentiation. | High on-off ratio in light and dark conditions. |

n-type semiconductor with good electron mobility. Together they form a bulk heterojunction (BHJ) structure (Fig. 10a), facilitating efficient electron-hole separation into free charge carriers upon light absorption [21]. The inclusion of interfacial layers such as PEDOT:PSS (anode) and Ca (cathode) in the photodiode facilitate transport and help align the energy levels between the electrodes and the active materials (Fig. 10b). Reducing the injection barriers results in a decrease of the threshold voltage. The choice of metal contacts has also been made to minimise the work function difference between the anode/cathode and the HUMO/LUMO levels of the active materials. Phototransistors require a high-quality dielectric to ensure a high capacitance per unit area, which can significantly lower the gate voltage required to turn on the device and induce the channel formation for charge transport. In this paper, 500nm of PMMA dielectric has been selected in order to increase capacitance (Fig. 11a).

#### *B. Electrical Properties and Photo Response - OghmaNano Simulation Results*

OghmaNano is a general purpose model for the simulation of opto-electronic devices [23] [24] [25]. The properties of both the photodiode and phototransistor were simulated in OghmaNano (Version 8.0.044) using finite element analysis (FEA) - see parameters in Table S4. The overall device geometry, including the thickness of the dielectric and semiconductor layers, is optimised to ensure that the electric field required to turn-on the device is minimised. For both devices, the substrate size in the simulations was set at $200\mu m \times 200\mu m$. Considering the current density output of both devices, selecting a substrate size of $200\mu m \times 200\mu m$ yielded an optimal current output. Additionally, this size is compatible with the eye's geometry and aligns with the optical analysis conducted by the developers of CurvIS [3].

Research shows that the minimum threshold light intensity for a human eye can be as little as $\approx 10^{-19}W/m^2$, however a more commonly reported value is between $10^{-12}$ to $10^{-10}W/m^2$ [26], which is equivalent to $\approx 10^{-13}Suns$ where $1Suns = 1000W/m^2$.

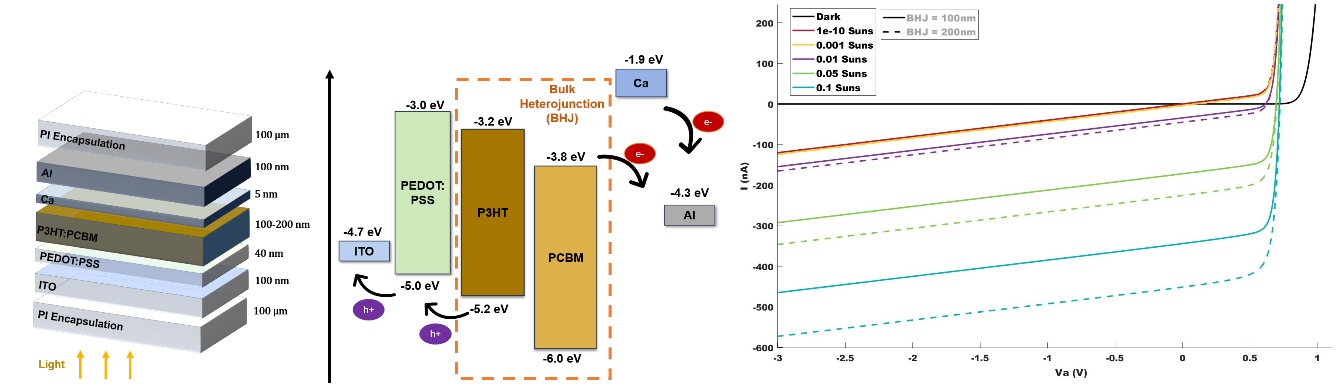


(a) (b) (c)

Figure 10: a) The semiconductor stack used in the photodiode simulations in OghmaNano. b) Energy Levels between contacts and active materials in the photodiode. Calcium is used as an electron transport layer, and PEDOT:PSS is used as a hole transport layer c) Output characteristics of a photodiode under increasing light intensity and different BHJ thicknesses. Photodiodes have a higher on/off ratio when operating in reverse bias. Note that $1 Suns = 1000 W/m^2$.

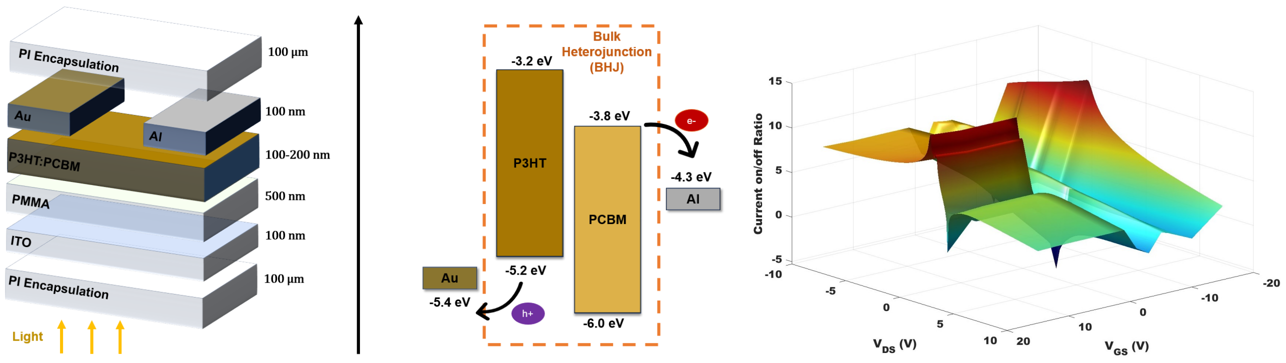


(a) (b) (c)

Figure 11: a) The semiconductor stack used in the phototransistor simulations in OghmaNano. ITO is used as the Gate in order to create the field effect and PMMA is the dielectric used. b) Energy Levels between contacts and active materials in the phototransistor. c) Surface plot to show a parameter sweep ($V_{GS}$ and $V_{DS}$), in order to optimise the on/off current ratio between Dark and $10^{-3} Suns$.

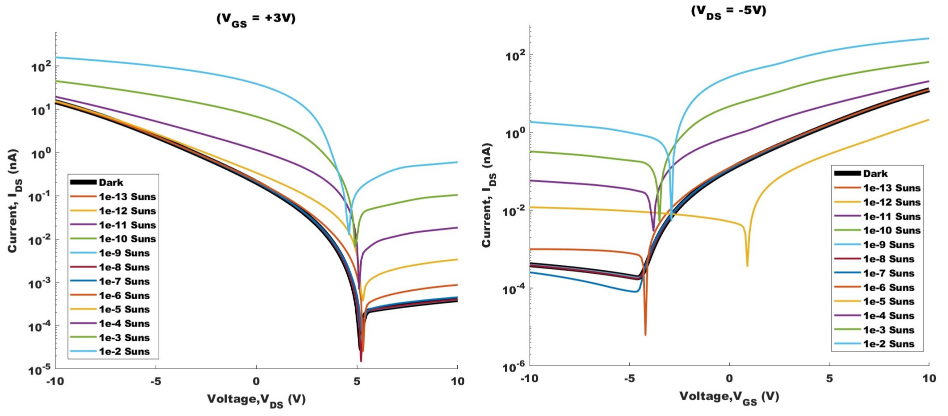


(a) (b)

Figure 12: a) Output characteristics of a phototransistor under increasing light intensity at $V_{GS} = 3V$. b) Transfer characteristics of a phototransistor under increasing light intensity at $V_{DS} = -5V$. Note that $1 Suns = 1000 W/m^2$.

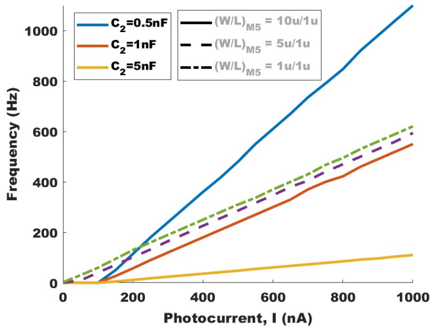


(a) Typical current for the photodiode response under light illumination between $1W/m^2 - 100W/m^2$.

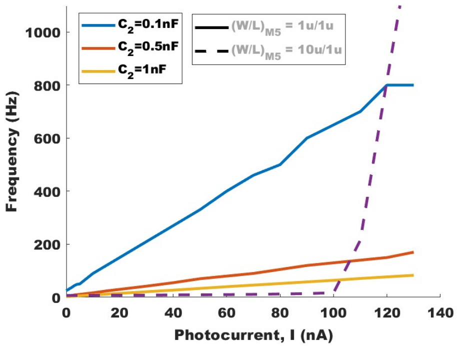


(b) Typical current for the phototransistor response under light illumination between $10^{-10}W/m^2 - 1W/m^2$.

Figure 13: Circuit Output frequency against photocurrent input, for parameter optimisation. M5 is the current sink that regulates the photo-voltage. C2 determines the pulse train frequency.
