## Supplementary Info Only for "An Integrated Photoreceptor-to-RGC Stimulation Circuit for Intraocular Visual Prostheses"

### Supplementary Information

Vedika Bedi supervised by Dr. Mujeeb U. Chaudhry

### I. NEURON Simulations

GitHub Link to the NEURON (Version 8.2) Code used in the simulations in this paper: https://github.com/VedikaBedi/NEURON_Code_VedikaBedi/tree/main

Table S1

Biophysical parameters used in NEURON 8.2 [1]. In NEURON, the RGC cell was modelled with; soma (D = 5 $\mu m$), a dendrite synapsing with a bipolar cell (D = 0.3 $\mu m$, L = 17 $\mu m$M), and the axon split into three parts consisting of the axon hillock (AH: D = 1.25 $\mu m$, L = 40 $\mu m$), Axon Initial segment, (AIS: D = 0.60 $\mu m$, L = 40 $\mu m$), and the remainder of the Axon (D=0.91$\mu m$, L= 1170 $\mu m$) [1].

| **Property** | **Value** |
|---|---|
| Axial Resistance | 110 Ωcm |
| Membrane Capacitance | 1 $\mu$F/cm$^2$ |
| ***Passive Membrane Model: Single passive membrane components to represent the overall passive (leak) conductance of the soma/dendrite membrane.*** | |
| Soma Passive Conductance | 0.001 S/cm$^2$ |
| **Soma Passive Reserve Potential - This is the Membrane Potential** | -70 mV |
| Dendrite Passive Conductance | 0.001 S/cm$^2$ |
| Dendrite Passive Reverse Potential | -65 mV |
| ***Hodgkin - Huxley (HH) Model: Conductance to do with the sodium, potassium, and leakage channels.*** | |
| Sodium ion channel Conductance | 0.12 S/cm$^2$ |
| Potassium ion channel Conductance | 0.036 S/cm$^2$ |
| Sodium ion Reverse Potential | 35 mV |
| Potassium ion Reverse Potential | -75 mV |
| Background Leakage Conductance to do with non-specific ion channels | 0.0003 S/cm$^2$ |
| Background Leakage Reverse Potential to do with non-specific ion channels. | -54.3 mV |

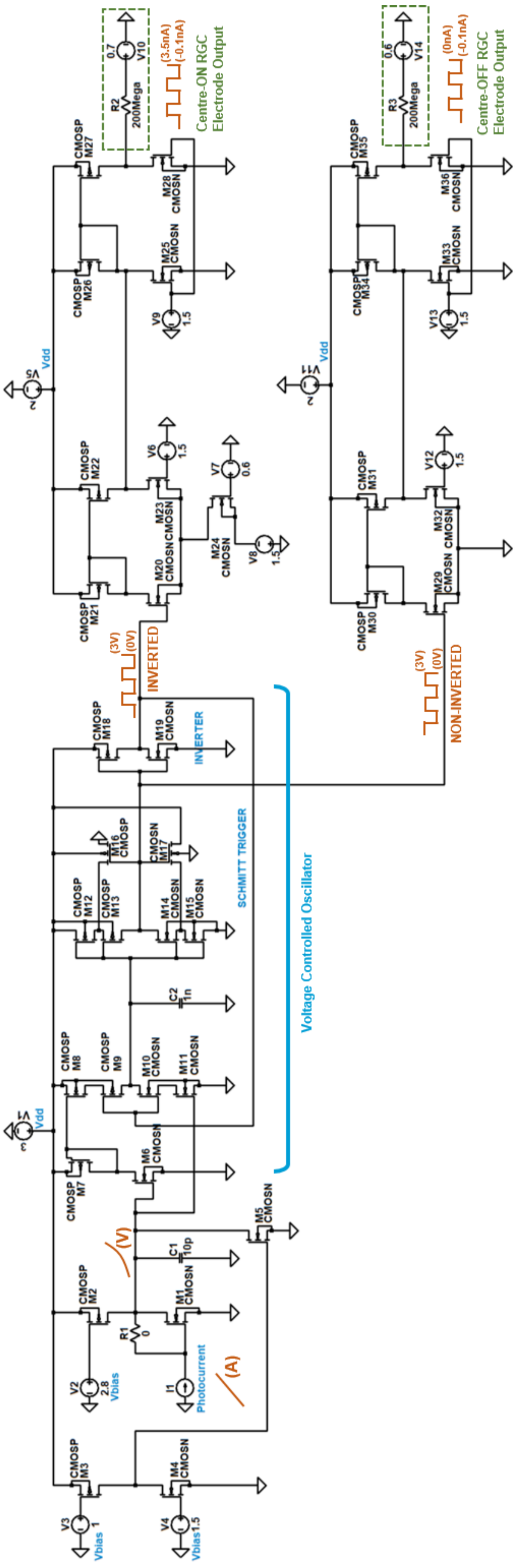


Figure S1: Proposed Circuit (enlarged figure)

### II. Circuit Modelling and LTSpice Simulations

Table S2
Breakdown of components values in the proposed neuron spiking circuit that converts a photocurrent to a suitable output current meeting the threshold requirements for RGCs.

| Component | Photodiode Circuit | Phototransistor Circuit |
|---|---|---|
| M1 – M4 | 10u/1u | 10u/1u |
| M5 | 5u/1u | 1u/1u |
| M7 – M24, M26 | 10u/1u | 10u/1u |
| M25 | 14u/1u | 14u/1u |
| M27-M29 , M31, M35 – M36 | 1u/1u | 1u/1u |
| M30, M32, M34 | 10u/1u | 10u/1u |
| M33 | 9u/1u | 9u/1u |
| C1 | 10pF | 10pF |
| C2 | 1nF | 0.1nF |
| Main Circuit $V_{dd}$ | 3V | 3V |
| RGC Centre-ON circuit $V_{dd}$ | 2 V | 2V |
| RGC Centre-ON circuit $V_{dd}$ | 2V | 2V |
| $V_{bias}$ M2 Gate Voltage | 2.8V | 2.8V |
| $V_{bias}$ M3 Gate Voltage | 1V | 1V |
| $V_{bias}$ M4 Gate Voltage | 1.5V | 1.5V |
| $V_{bias}$ M23 Gate Voltage | 1.5V | 1.5V |
| $V_{bias}$ M24 Gate Voltage | 0.6V | 0.6V |
| $V_{bias}$ M24 Source Voltage | 1.5V | 1.5V |
| $V_{bias}$ M25 Gate Voltage | 1.5V | 1.5V |
| $V_{bias}$ M32 Gate Voltage | 1.5V | 1.5V |
| $V_{bias}$ M33 Gate Voltage | 1.5V | 1.5V |
| R2 and R3 Electrodes | 200M Ohms | 200M Ohms |
| R2 Electrode Return Voltage | 0.7V | 0.7V |
| R3 Electrode Return Voltage | 0.6V | 0.6V |

Table S3
MOSFET parameters used in the Level 3 model in LTSpice (Version 24.0.9).

| LTSpice Parameters | Ideal NMOS | Ideal PMOS |
|---|---|---|
| Oxide Thickness,$T_{ox}$ | 200e-10 | 200e-10 |
| Substrate Doping, $NSUB$ | 1e17 | 1e17 |
| Body Effect Parameter, $\gamma$ | 0.5 | 0.6 |
| Surface Potential, $\phi$ | 0.7 | 0.7 |
| Threshold Voltage,$V_{th}$ | 0.5 | -0.5 |
| Mobility, $\mu$ | 650 | 250 |
| Drain induced barrier lowering, $ETA$ | 3.0e-6 | 0 |
| Transconductance Parameter, $K_P$ | 120e-6 | 40e-6 |
| Sheet Resistance, $R_{SH}$ | 0 | 0 |
| Junction Depth, $XJ$ | 500e-9 | 500e-9 |
| Gate drain overlap capacitance per unit width, $C_{GDO}$ | 200e-12 | 200e-12 |
| Zero-bias bulk junction capacitance, $C_J$ | 400e-6 | 400e-6 |
| Zero-bias sidewall junction capacitance, $C_{JSW}$ | 300e-12 | 300e-12 |
| Maximum drift velocity of carriers, $V_{Max}$ | 1e5 | 5e4 |
| Width effect on threshold voltage, $\Delta$ | 3.0 | 0.1 |
| Mobility reduction coefficient, $\theta$ | 0.1 | 0.1 |
| Saturation field factor,$\kappa$ | 0.3 | 1 |
| Fast surface state density, $NFS$ | 1e12 | 1e12 |
| Lateral diffusion, $LD$ | 100e-9 | 100e-9 |
| Type of gate material relative to the substrate,$TPG$ | 1 | -1 |
| Gate-source overlap capacitance per unit width, $C_{GSO}$ | 200e-12 | 200e-12 |
| Gate-bulk overlap capacitance per unit length,$C_{GBO}$ | 1e-10 | 1e-10 |
| Bulk junction potential, $PB$ | 1 | 1 |
| Bulk junction grading coefficient, $MJ$ | 0.5 | 0.5 |
| Sidewall junction grading coefficient, $MJSW$ | 0.5 | 0.5 |

```
.MODEL CMOSN NMOS LEVEL  = 3
+ TOX    = 200E-10        NSUB   = 1E17          GAMMA  = 0.5
+ PHI    = 0.7            VTO    = 0.5          DELTA  = 3.0
+ UO     = 650            ETA    = 3.0E-6        THETA  = 0.1
+ KP     = 120E-6         VMAX   = 1E5           KAPPA  = 0.3
+ RSH    = 0              NFS    = 1E12          TPG    = 1
+ XJ     = 500E-9         LD     = 100E-9
+ CGDO   = 200E-12        CGSO   = 200E-12       CGBO   = 1E-10
+ CJ     = 400E-6         PB     = 1             MJ     = 0.5
+ CJSW   = 300E-12        MJSW   = 0.5

.MODEL CMOSP PMOS LEVEL  = 3
+ TOX    = 200E-10        NSUB   = 1E17          GAMMA  = 0.6
+ PHI    = 0.7            VTO    = -0.5         DELTA  = 0.1
+ UO     = 250            ETA    = 0             THETA  = 0.1
+ KP     = 40E-6          VMAX   = 5E4           KAPPA  = 1
+ RSH    = 0              NFS    = 1E12          TPG    = -1
+ XJ     = 500E-9         LD     = 100E-9
+ CGDO   = 200E-12        CGSO   = 200E-12       CGBO   = 1E-10
+ CJ     = 400E-6         PB     = 1             MJ     = 0.5
+ CJSW   = 300E-12        MJSW   = 0.5
```

Figure S2: .MODEL statements in LTSpice to set up the Level 3 MOSFET model

### III. OGHMANANO SIMULATIONS

Table S4
OGHMANANO (VERSION 8.0.044) SIMULATION PARAMETERS - ELECTRICAL PARAMETERS OF THE ACTIVE MATERIAL (P3HT:PCBM)

| OghmaNano Electrical Parameters | Value |
|---|---|
| Free Carriers | |
| Electron Mobility | 2.48e-07 $m^2V^{-1}s^{-1}$ - Symmetric |
| Hole Mobility | 2.48e-07 $m^2V^{-1}s^{-1}$ - Symmetric |
| Effective Density of free electron states (@300K) | 1.28e27 $m^{-3}$ |
| Effective Density of free hole states (@300K) | 2.86e25 $m^{-3}$ |
| $n_{free}$ to $p_{free}$ Recombination rate constant | $0.0m^3s^{-1}$ |
| Free carrier statistics | Maxwell Boltzmann - Analytic (photodiode)<br>Fermi-Dirac Numerical (phototransistor) |
| Non-equilibrium SRH traps | |
| DoS Distribution | Exponential |
| Electron trap density | 3.8e26 $m^{-3}eV^{-1}$ |
| Hole trap density | 1.45e25 $m^{-3}eV^{-1}$ |
| Electron tail slope | 0.04 eV |
| Hole tail slope | 0.06 eV |
| Free electron to Trapped electron | 2.5e-20 $m^{-2}$ |
| Trapped electron to Free hole | 1.32e-22 $m^{-2}$ |
| Trapped hole to Free electron | 4.67e-26 $m^{-2}$ |
| Free hole to Trapped hole | 4.86e-22 $m^{-2}$ |
| Number of traps | 20 bands |
| Electrostatics | |
| Xi | 3.8 eV |
| Eg | 1.3 eV |
| Relative Permittivity | 3.8 au |

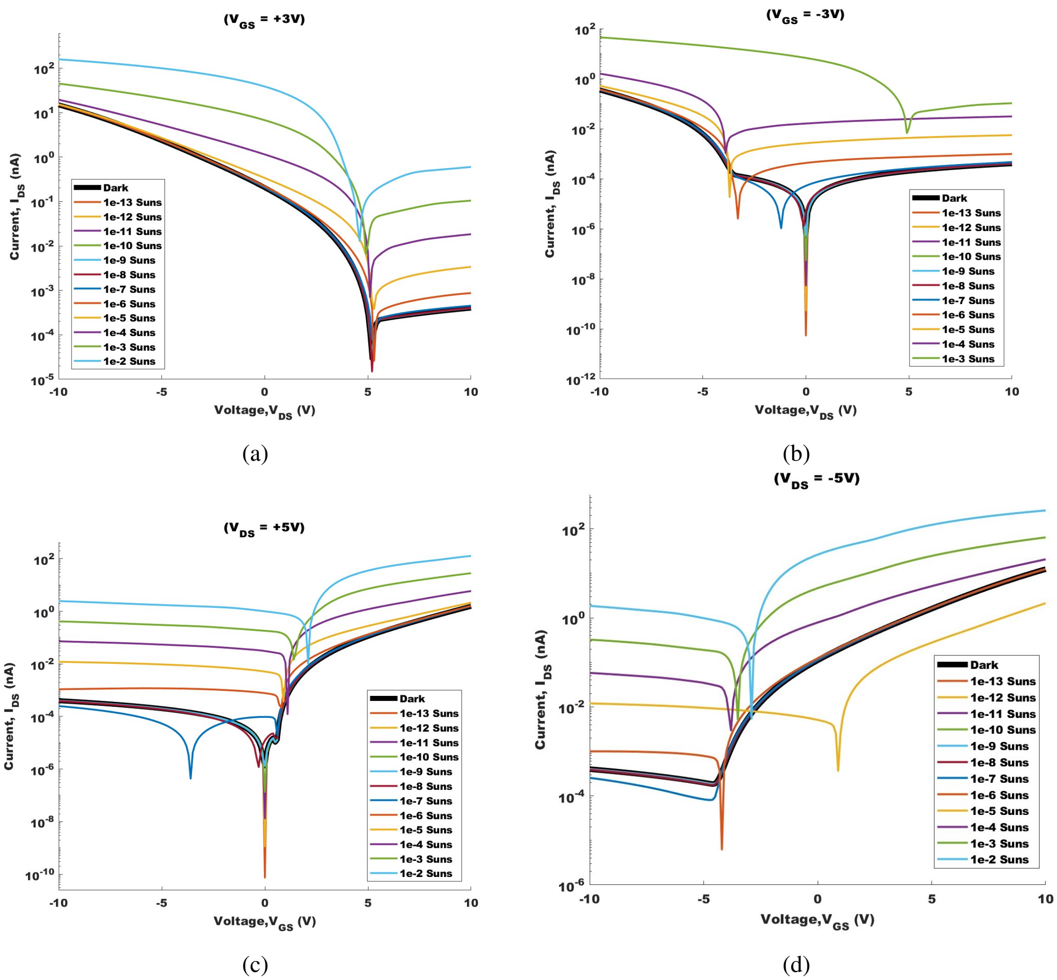


Figure S3: Output and Transfer Characteristics of the phototransistor. This provides better illustration of how the choice of positive and negative voltages impacts the characteristics of the phototransistor. Note this is due to an unequal blend of P3HT:PCBM being present in the semiconductor. This material database has been taken from a solar cell model, and so the concentration blend is yet to be optimised.

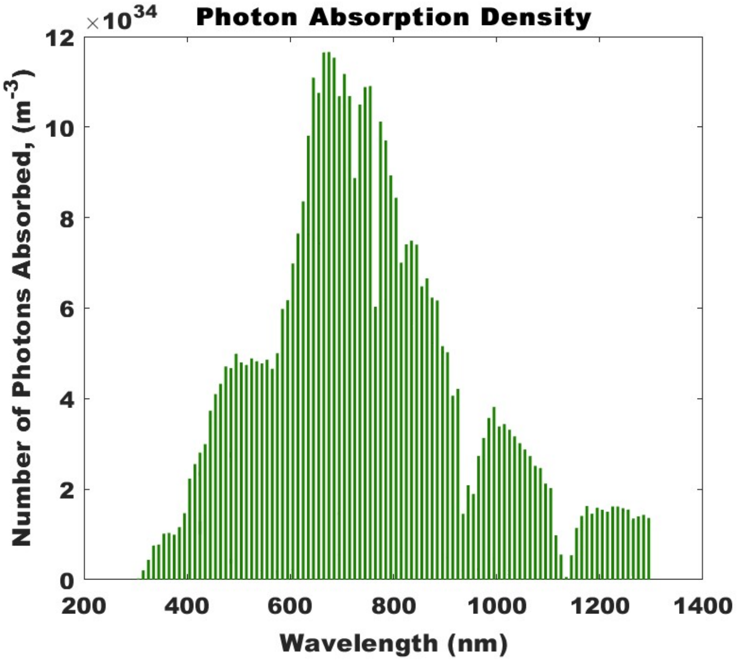


Figure S4: The P3HT:PCBM blend has excellent photon absorptivity at all visible light wavelengths, with the highest absorption at 650nm. The photoreceptors will however, require an Infrared filter as it also is shown to have good absorption between 700-1200nm.

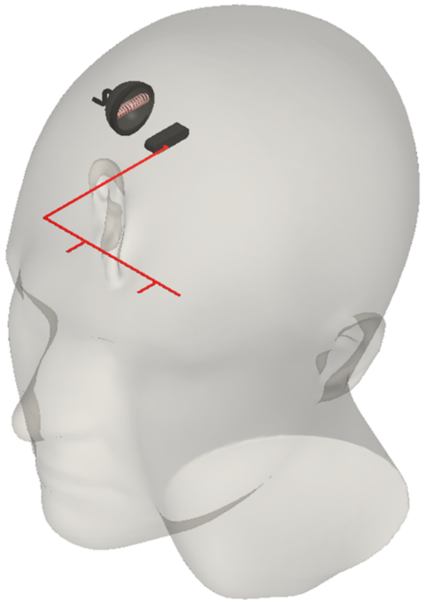

Figure S5: Not to Scale CAD image of a human head fitted with an inductive coupling device with power wires for each eye.
